# Exploration of T-cell immune responses by expression of a dominant-negative SHP1 and SHP2

**DOI:** 10.1101/2023.04.26.538403

**Authors:** Julia Taylor, Anna Bulek, Isaac Gannon, Mathew Robson, Evangelia Kokalaki, Thomas Grothier, Callum McKenzie, Mohamed El-Kholy, Maria Stavrou, Charlotte Traynor-White, Wen-Chean Lim, Panagiota Panagiotou, Saket Srivastava, Vania Baldan, James Silibourne, Mathieu Ferrari, Martin Pule, Simon Thomas

## Abstract

SHP1 and SHP2 are SH2 domain-containing proteins which have inhibitory phosphatase activity when recruited to phosphorylated ITIMs and ITSMs on inhibitory immune receptors. Consequently, SHP1 and SHP2 are key proteins in the transmission of inhibitory signals within T cells, constituting an important point of convergence for diverse inhibitory receptors. Therefore, SHP1 and SHP2 inhibition may represent a strategy for preventing immunosuppression of T cells mediated by cancers hence improving immunotherapies directed against these malignancies. Both SHP1 and SHP2 contain dual SH2 domains responsible for localization to the endodomain of inhibitory receptors and a protein tyrosine phosphatase domain which dephosphorylates and thus inhibits key mediators of T cell activation. We explored the interaction of the isolated SH2 domains of SHP1 and SHP2 to inhibitory motifs from PD1 and identified strong binding of both SH2 domains from SHP2 and more moderate binding in the case of SHP1. We next explored whether these a truncated form of SHP1/2 comprising only of SH2 domains (dSHP1/2) could act in a dominant negative fashion by preventing docking of the wild type proteins. When co-expressed with CARs we found that dSHP2 but not dSHP1 could alleviate immunosuppression mediated by PD1. We next explored the capacity of dSHP2 to bind with other inhibitory receptors and observed several potential interactions. In vivo we observed that the expression of PDL1 on tumor cells impaired the ability of CAR T cells to mediate tumor rejection and this effect was partially reversed by the co-expression of dSHP2 albeit at the cost of reduced CAR T cell proliferation. Modulation of SHP1 and SHP2 activity in engineered T cells through the expression of these truncated variants may enhance T cell activity and hence efficacy in the context of cancer immunotherapy.

## Introduction

T cell activation is regulated through a dynamic balance between tyrosine phosphorylation and dephosphorylation through the interaction of complementary kinases and phosphatases with membrane bound receptors. The initial signal for T cell activation is the phosphorylation of immunoreceptor tyrosine activation motifs (ITAMs) of the T cell receptor – (TCR)-CD3 complex. Upon antigen recognition, an immunological synapse forms, from which CD45 phosphatases are excluded in the favor of Lck and related kinases, pushing the balance towards phosphorylation and hence activation in a process referred to as kinetic segregation^1^. A further input comes from phosphatase activity of two related SH2 domain-containing phosphatases: SHP1 and SHP2^2^. These phosphatases are recruited to phosphorylated immunoreceptor inhibitory motifs (ITIMs) and immunoreceptor tyrosine switch motifs (ITSMs) on inhibitory immune receptor endodomains^3^ and, as many of these inhibitory receptors signal through SHP1 and SHP2, these phosphatases constitute a key convergence or “pinch point” in immune cell inhibitory signaling.

A key factor preventing the successful rejection of tumor by immune effector cells is the immune microenvironment tipping the balance in favor of T cell inhibition and inactivity. Endogenous tumor specific T cells and adoptively transferred immune effectors such as TILs, transgenic TCR and CAR T cells can be inhibited by a range of inhibitory microenvironmental signals emanating from tumor or associated inhibitory cells. Inhibitory factors include adenosine, TGFβ, IL10 and indoleamine 2, 3- dioxygenase and others^4^. Additionally inhibitory ligands expressed on the surface of tumor and inhibitory cells can interact with ITIM/ITSM-containing receptors on T cells to mediate their inhibition. Chief among these is PD1, however ITIM and ITSM motifs have been identified in a variety of other inhibitory receptors expressed by T cells such as BTLA^5^, TIGIT^6^ and 2B4^7^, suggesting that SHP1 and SHP2 may therefore represent a potential target for intervention for the relief of all such inhibition.

Several approaches have been described to block immune suppression in the tumor microenvironment. The simplest approach is administration of blocking antibodies. Adoptively transferred engineered T-cells can be co-administered with blocking antibodies, but ex-vivo manipulation affords approaches restricted to the engineered cells. This includes gene-editing of inhibitory receptors^8^, or expression of switch receptors^9^, or engineered secretion of blocking antibody fragments^10^. Considering the position of SHP1 and SHP2 at the crux of multiple inhibitory pathways, we considered that discretely blocking these proteins in the context of an engineered T-cell, was as strategy worthy of exploration with possible advantages over approaches directed at individual receptors including the possibility of blocking multiple inhibitory signals.

SHP1 and SHP2 share a similar overall structure possessing two N-terminal src homology 2 (SH2) domains, a catalytic protein tyrosine phosphatase (PTP) domain and a C-terminal tail^2^. We hypothesized that a truncated, catalytically inactive form of SHP1/2 comprising only of the SH2 domains would compete for phospho-ITIM/ITSM sequences, preventing recruitment of wild type SHP1/2. We first explored the binding of SHP1 and SHP2 to unphosphorylated and phosphorylated ITIM and ITSM motif from PD1, as well as motifs derived from a range of other inhibitory receptors including CTLA4, BTLA, TIGIT and 2B4. We next explored the consequence of inhibiting SHP1 and SHP2 through expression of truncated forms in the context of CAR T cells, investigating the phenotypic, functional and transcriptional consequences of expressing a truncated SHP2 and comparing its effect on PD1-mediated inhibition of CAR T cells in vitro with PD1 blockade using pembrolizumab. Finally, we demonstrated the function of truncated SHP2 *in vivo* in the context of a CD19 CAR and PD1/PDL1.

## Materials and Methods

### Cell lines

HEK-293T (ATCC; ATCC®CRL-11268™) were cultured in Iscove’s modified Dulbecco’s medium (IMDM) (Lonza, Basel, Switzerland) supplemented with 10% FBS (Labtech, Heathfield, UK) and 5 mM GlutaMAX (Invitrogen, CA). SupT1 (ECACC; 95013123), NALM6 (DSMZ; ACC 128) lines were cultured in complete RPMI (RPMI-1640, Lonza, Basel, Switzerland) supplemented with 10% FBS and 2 mM GlutaMAX (Gibco; 35050061).

### DNA Construct generation

All open reading frames were cloned into the MoMLV-based retroviral genome construct SFG. Linear DNA fragments (gBlocks), encoding codon optimised open reading frames (GeneArt), were synthesised (IDT) and amplified using Q5 DNA polymerase (NEB) and oligonucleotides (IDT). The resulting PCR products were fractionated on an agarose gel, purified by gel extraction (Qiagen), and digested with restriction endonucleases (NEB). Digested DNA fragments were purified (Qiagen) and ligated to gel-purified plasmid backbones using T4 DNA ligase (Roche). Competent DH5a *E. coli* (NEB) were transformed with the ligation reactions, plated onto LB agar containing ampicillin (final concentration of 100 mg/mL) and incubated overnight at 37oC.

### Recombinant protein (SH2 domain) production and purification

DNA constructs encoding SHP1 or SHP2 SH2 domains were cloned into pGMT7 expression vector, transformed into *E. coli* BL21(DE3)-pLysS (Novagen), expressed as inclusion bodies and refolded using previously described methods^11^. Purification of histidine tagged proteins was performed using FPLC chromatography on HiTrap TALON crude 5 ml column (Cytiva) according to manufacturer’s recommendations. The sample was further purified using size exclusion chromatography on a HiLoad 16/600 Superdex 75 column (Cytiva). The final sample was validated using SDS-PAGE and thermostability assay (Nanotemper NT.48 nanoDSF).

### Phosphopeptide ELISA

Phosphorylated and unphosphorylated peptides were synthesised by Genscript. Peptides were coated at 0.5 µg/ml on Nunc MaxiSorp 96 well plates overnight at RT. Plates were washed 3 times in PBS 0.05% Tween20 and blocked in PBS 2% BSA for 1 h at RT. Plates were then washed 3 times in PBS 0.05% Tween20 and incubated with 50 µl of test protein at 10 µg/ml with 1:3 serial dilutions in PBS 0.5% BSA. Plates were washed 3 times in PBS 0.05% Tween20 and incubated with secondary antibody (Anti-His-Tag Antibody (H-3): sc-8036 from Santa Cruz) at 1 µg/ml in PBS 0.5% BSA for 1 h at RT. Plates were then washed 3 times in PBS 0.05% Tween20 and incubated with 45 µg/well of Ultra-TMB reagent (Thermo Scientific) before blocking with 45 µl of 1M H2SO4. Absorbable was read at 450nm.

### Surface Plasmon Resonance (Biacore)

The Surface Plasmon Resonance (SPR) experiments were performed using a BIAcore^TM^ T200 (GE Healthcare) equipped with Series S Sensor Chip CAP (Cytiva). Biotinylated Ligands were immobilized at a density of 15 RU under a flow rate of 10 ul/min in HBS-EP+ buffer. To collect kinetic binding data, analytes in HBS-EP^+^ buffer were injected over four flow cells at concentrations of 250 nM, with 2-fold serial dilutions, at a flow rate 30 µl/min and at a temperature of 25^0^C. The complex was allowed to associate and dissociate for 150 and 300s, respectively. To remove Biotin CAPture Reagent as well as the ligand and any bound analyte, surfaces were regenerated with 120s injections of regeneration solution at a flow rate 30 ul/min. The data were fit to a 1:1 Langmuir interaction model.

### Flow Cytometry

Flow cytometry was performed using MACSQuant 10 and X flow cytometers (Miltenyi Biotec, Germany). All labelling was carried out at room temperature (RT) for 10mins protected from light with antibodies diluted in cell staining buffer (Biolegend; 420201). For AnnexinV staining, cells were harvested and stained for CD3 and RQR8 expression, washed once in PBS, washed once in Annexin V buffer and resuspended in Annexin V binding buffer with Annexin V-BV421 (Biolegend; 640924) and incubated for 15 minutes at room temperature protected from the light. Cells were then washed and resuspended in Annexin V binding buffer containing 7-AAD and analysed by flow cytometry. Apoptotic cells were defined as being Annexin V^+^ /7-AAD^−^.

Antibodies used were: CCR7-PE (Miltenyi Biotec; 130-119-583), CD19–APC Cy7 (BioLegend; 302218), CD25-BV421 (Biolegend; 356114), CD27-VioBright515 (Miltenyi Biotec; 130-120-028), CD3–PE Cy7 (BioLegend; 344816), CD3-VioGreen (Miltenyi Biotec; 130-113-142), CD45RA- APCVio770 (Miltenyi Biotec; 130-117-747), CD69-FITC (Biolegend; 310904), CD8-APC-Cy7 (Biolegend; 301016), CD8-Vioblue (Miltenyi Biotec; 130-110-683), CD95-PEVio770 (Miltenyi Biotec 130-113-006), HA–AF488 (Biolegend; 901509), KLRG1-APC-Vio77 (Miltenyi Biotec; 130-120-423), LAG3-VioBright515 (Miltenyi Biotec; 130-120-012), PD1-PE (Miltenyi Biotec; 130-120- 382), CD34 (RQR8)-PE (R&D Systems; FAB7227P), CD34 (RQR8)-APC (R&D Systems; FAB7227A), SYTOX™ AADvanced™ Dead Cell Stain (ThermoFisher; S10274), TIM3-PEVio770 (Miltenyi Biotec; 130-121-334), GD2-APC (Biolegend; 357306), HVEM-PE (Biolegend; 318806), PDL1-PE (Biolegend; 329706), CD48-PE (Biolegend; 336708), CD80-FITC (Biolegend; 305206), CD155-PE (Biolegend; 337610). Anti-CD19 CAR was detected using an anti-idiotype antibody specific for the HD37 aCD19 binder (in-house) and Goat anti-Mouse IgG2a secondary – AF647 (Invitrogen; A21241).

### Retroviral production

HEK-293T cells (1.5x10^6^) were transiently transfected with an RD114 envelope expression plasmid (RDF, a gift from M. Collins, University College London), and a Gag-pol expression plasmid (PeqPam-env, a gift from E. Vanin, Baylor College of Medicine), and transgene expressed in a retroviral (SFG) vector plasmid at a ratio of 1:1.5:1.5 (total DNA=12.5ug). Transfections were carried out with GeneJuice (Millipore; 70967-4) according to the manufacturer’s guidelines and viral supernatants were harvested 48 hours post transfection and stored at -80°C.

### Transduction of cells

Peripheral Blood Mononuclear Cells (PBMCs) were isolated from whole blood by density centrifugation via Ficoll-Paque PLUS (GE-Healthcare; GE17-1440-03) according to manufacturer’s protocol. Isolated PBMCs were activated using 50ng/ml of aCD3 (Miltenyi Biotec; 130-093-387)/ aCD28 (Miltenyi Biotec; 130-093-375) followed by 50U/ml of IL2 (Miltenyi Biotec; 130-097-743) after 24 hours. 48hrs post activation, 1x10^6^ PBMCs were plated on retronectin (Takara Clonetech; T100B) on 6 well plates (Corning; 351146) with retrovirus vector and spun for 1000xg 40mins at RT. Tumor cell lines were transduced with retroviral supernatants on retronectin-coated 6-well plates at a concentration of 0.25x10^6^ cells/ml. Expression of the transgene was assessed by flow cytometry 72 hours post transduction. In order to generate cells expressing PDL1 and/or GD2, cell lines were transduced with retroviral constructs encoding PDL1 and/or the biosynthetic enzymes GD3 synthase and GD2 synthase linked by a viral 2A sequence. Expression of these two synthases results in the synthesis and expression on the cell surface of GD2^12^. Transduced cells expressing PDL1 and/or GD2 were isolated by FACS.

### Cytotoxicity assay

Transduced T cells were depleted of CD56 cells using the EasySep Human CD56 positive selection kit (Stemcell Technologies; 17855) and rested for 24 hours. Transduced cells were co-cultured with target cells at the indicated effector : target (E:T) ratios from with target cells remaining constant at the indicated number of target cells/co-culture. At the end of the co-culture period target cells were enumerated by flow cytometry and cytotoxicity calculated as a percentage of the number of target cells recovered from co-cultures with non-transduced (NT) T cells.

### Cytokine ELISAs

Cytokine concentrations in supernatants harvested from CAR T cell: target cell co-cultures were determined by ELISA utilizing IFNγ and IL2 Deluxe kits (Biolegend; 431806 and 430106) according to the manufacturer’s instructions.

### Proliferation assays

PBMCs were labelled with Cell Trace Violet (CTV) according to manufacturer’s instructions (ThermoFisher Scientific; C34557) and cultured with 0.5x10^5^ target cells at 1:1 E:T ratio for 96h. Numbers of CD3^+^ RQR8+ T cells and CTV MFI were assessed by flow cytometry.

### Serial rechallenge assays

For serial rechallenge assays to measure proliferation and activation-induced cell death, T cells were stimulated with 5x10^4^ of the indicated target cells at an E:T ratio of 1:1 or 1:4 and incubated at 37°C/5% CO2 for 72 hours. Co-cultures were harvested and T cells enumerated and assessed for apoptosis as described above prior to restimulation with 5x10^4^ fresh target cells as before. In order to measure cytotoxicity 1x10^5^ Skov3 cells engineered to express GD2 alone or in combination with PDL1 and the fluorescent protein mKate were plated in 24-well cell culture plates and allowed to adhere for 24 hours prior to the addition of 4x10^5^ CAR T cells to achieve an effector:target ratio of 4:1. Target cell killing was then monitored for 72 hours using an Incucyte Live Cell Imager by enumerating the number of fluorescent Skov3 cells remaining at hourly intervals. T cells were then harvested and rested for 72 hours prior to initiation of a second co-culture with Skov3 target cells as before.

### Plate bound CAR Assays

Nunc MaxiSorp 96 well plates were coated overnight at 4°C with increasing concentrations of anti- idiotype antibody specific to the binding domain of the anti-CD19 CAR. T cells transduced to express either the anti-CD19 CAR alone or alongside dSHP2 were stimulated on the plates for 24 or 72 hours prior to being harvested and transduced (CD3+/RQR8+) T cells were assessed for the expression of markers of activation (CD69, CD25), memory (CD45RA, CD95, CD27, CCR7) or exhaustion (PD1, KLRG1, TIM3, Lag3) by flow cytometry.

### Transcriptomic analysis

T cells expressing the CAR alone or in combination with the dSHP2 module were stimulated for 24 hours using 5µg/ml anti-CAR idiotype antibody coated onto Nunc MaxiSorp 96 well plates. RNA was extracted from CAR T cells using a Qiagen RNeasy Mini Kit (Qiagen; #74104) and RNA was quantified by NanoDrop Spectrophotometer (ThermoFisher). The quality of the RNA was assessed with Agilent TapeStation (Agilent, RRID:SCR_019547). RNA sequencing was carried out on the NanoString nCounter SPRINT according to manufacturer instructions and using 80ng of RNA per sample.

### SHP2 gene disruption

Expression of the SHP2 (PTPN11) gene was disrupted by Cas9-mediated indel formation. For each sgRNA, 2 samples of 5x106 PBMCs were nucleofected in 100 mL of P3 buffer (Lonza; V4XP-3024) with 125 pmol of Alt-R Sp HiFi Cas 9 nuclease v3 (IDT; 108210621) complexed with 187.5 pmol of the PTPN11 sgRNA (GAGACUUCACACUUUCCGUU) or a non-targeting sgRNA (GCACUACCAGAGCUAACUCA; Synthego) using pulse code EH-115. Complexing of Cas9 and the sgRNA was carried out at ambient temperature for 10 minutes prior to nucleofection. Following nucleofection, PBMCs were allowed to recover for 10 minutes at 37oC/5% CO2 before the addition of 400 mL complete RPMI medium transfer to a single well of a 6-well plate containing 2 mL of complete RPMI medium supplemented with 50 U/mL of IL-2. PBMCs were transduced with retroviral constructs 24 hrs after nucleofection. Samples PBMCs were collected for genotyping, Western blotting and functional assessment 6 days after nucleofection.

### Estimation of SHP2 gene disruption by inference of CRISPR edits (ICE)

Genomic DNA was extracted from PBMCs using the GenElute Blood Genomic DNA Kit (Sigma; NA2010) according to the manufacturer’s instructions. The sequence flanking the SHP2 sgRNA target site was amplified by PCR from 100 ng of genomic DNA using Q5 DNA polymerase (NEB; M0491L) and the primers, SHP2_forward 5’-AGGGACAGGGAAGGTCTTGA-3’ and SHP2_reverse 5’-AAACTCGAAATGCAGGCAGC-3’ (IDT). The PCR products were Sanger sequenced (Source Bioscience) and indel formation and knockout efficiencies determined by ICE (Synthego), comparing SHP2-edited and non-target-edited sequences to an unedited control.

### Western blotting and intracellular phospho-protein staining

Cells were stimulated as indicated then 0.5x10^6^ cells were harvested and pelleted at 400xg for 2 minutes. The pellets were washed with ice-cold PBS, centrifuged as before and lysed in 50µL of 1x RIPA buffer (Millipore) supplemented with 1x protease and phosphatase inhibitor (Abcam). The cells were incubated on ice for 15 minutes, vortexed and incubated for further 15 minutes. The lysate was clarified by centrifuging the cells at 13000xg for 10 minutes at 4°C, protein concentration was determined using the BCA Protein assay Kit (Pierce) and stored at -80°C until use. Prior to gel electrophoresis lysates were mixed with NuPAGE™ LDS Sample Buffer (4X) (Invitrogen™) and heated at 95°C for 10 minutes. Samples (4µg) were resolved on a premade 4–20% Mini- PROTEAN® TGX™ Precast Gel (Abcam).

Post-separation, SDS-PAGE gel was transferred to Trans-Blot Turbo Midi 0.2 µm PVDF membrane (Abcam) using Trans-Blot® Turbo™ Midi by semi-dry transfer (Abcam). Following transfer, membranes were blocked with 1x TBST (Sigma) supplemented with 5% BSA for 24 hours at 4°C. Membrane staining was undertaken with primary antibody diluted to appropriate concentration in 1x TBST supplemented with 5% BSA for 1 hour on an orbital shaker at 480rpm at RT. The sample was washed 3 times with 1x TBST for 10 minutes each time and secondary staining was performed as per the primary stain. Following the last TBST wash, the membrane was washed with distilled water, then the membrane developed by incubation with Pierce ECL Plus Western Blotting Substrate (Abcam) or Immobilon Western Chemiluminescent HRP substrate (Millipore), according to the manufacturer’s instructions.

Antibodies used for Western blotting were as follows: phosphorylated ERK (Cell Signaling technology; 9101), ERK (Cell Signaling technology; 4695), SHP2 (BioLegend; 948202), GAPDH (Cell Signaling technology; 2118), Anti-Rabbit IgG HRP (Cell Signaling technology; 7074) or phosphorylated ribosomal protein S6 analysis, cells were stimulated as indicated then harvested by centrifugation at 400xg for 2 minutes. Cells were stained with Fixable Viability Dye eFluor™ 780 (65-0865-14) and CD34 (RQR8)-APC (R&D Systems; FAB7227A), washed in PBS and resuspended in BD Cytofix/Cytoperm™ (BD; 51-2090KZ). Cells were washed in BD Perm/Wash™ buffer (BD;51-2091KZ) and stained with anti Phospho-S6 Ribosomal Protein (Ser240/244)-PE (Cell Signaling technology; 14236) diluted in the BD Perm/Wash Buffer and analysed by flow cytometry.

### In vivo studies

All animal studies were performed under a UK Home Office–approved project license. Female, 6– 10-week-old NSG mice (Charles River Laboratory) were raised under pathogen-free conditions.

0.5 x10^6^ Nalm6 WT or PDL1-Nalm6, engineered to express firefly luciferase and a HA tag, were inoculated intravenously into NSG mice on day -4, respective to CAR T infusion. Tumor engraftment was measured by bioluminescent imaging utilizing the IVIS spectrum system (PerkinElmer, Waltham, MA) after intra peritoneal injection of luciferin. Mice were randomized on day -1 and the following day 1x10^6^ CAR T-cells were injected intravenously. Tumor growth was monitored by bioluminescent imaging. On day 14 post CAR T injection half the mice in each cohort were random selected for culling and CAR T cells in the blood enumerated by flow cytometry.

### Statistical analysis

All statistics was generated using GraphPad Prism, analysis was performed using two-way ANOVA. Significance is defined as: NS, not significant; *P < 0.05; **P < 0.01; ***P < 0.001; ****P < 0.0001

## Results

### SH2 domains from SHP2 selectively bind phospho-ITIM/ITSMs from PD1

The binding of SHP proteins to PD1 results in the dephosphorylation of key mediators of CAR T cell activation (Figure 1A). We first sought to determine the specificity of SHP1 and SHP2 SH2 domains to the PD1 ITIM and ITSM. The N and C terminal SH2 domains from SHP1 and SHP2 respectively were expressed as individual domains or linked in tandem and purified for subsequent experiments. Phosphorylated and unphosphorylated peptides corresponding to the PD1 ITIM and ITSM motifs were synthesized chemically (Supplementary table 1) and the interaction between SH2 domains and peptides were measured by ELISA and Biacore (Figure 1B and C).

**Figure 1:**
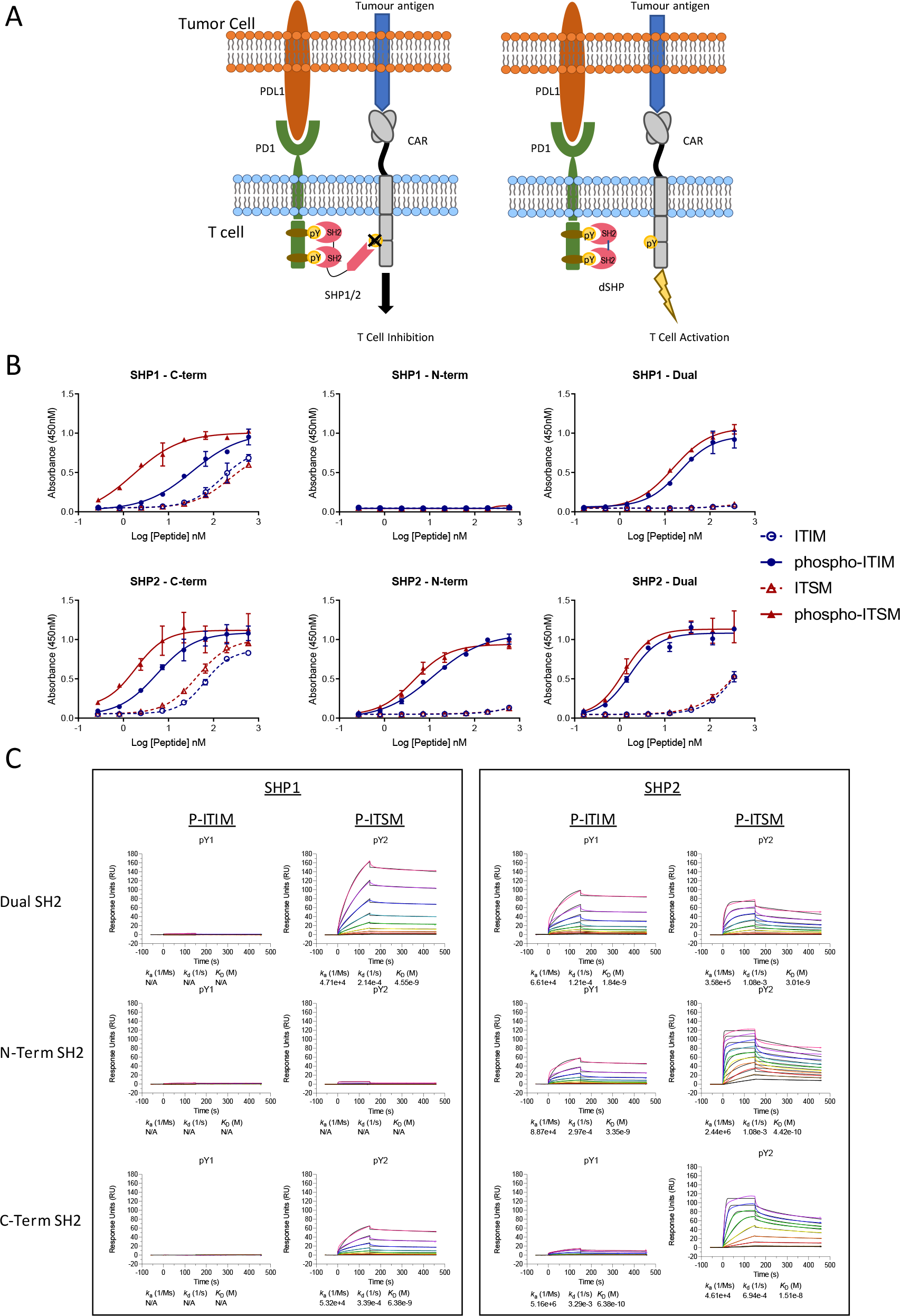
**Binding of the SH2 domains of SHP1 and SHP2 to the ITIM and ITSM sequences of PD1. A**) Inhibition of SHP activity by the isolated SH2 domains lacking catalytic activity. The truncated SH2 domains of SHP1 and SHP2, lacking the phosphatase domain, occupy the phosphorylated endodomain of PD1 and prevent native SHP1 and 2 from binding and inhibiting CAR function. **B**) Binding of the SH2 domains of SHP1 and SHP2 to the phosphorylated ITIM and ITSM peptides of PD1 by ELISA. Plates were coated with peptide and the indicated concentration and the purified, His-tagged SH2 domains of SHP1 or SHP2 were allowed to bind overnight. Binding of the indicated SH2 domain was measured through detection of the His tag and EC50 values for the phosphorylated peptides calculated. **C**) Binding of the SH2 domains of SHP1 and SHP2 to the phosphorylated ITIM and ITSM peptides of PD1 by Surface Plasmon resonance. Peptides corresponding to the phosphorylated or non-phosphorylated PD1 ITIM and ITSM motifs were immobilized.

The dual SH2 domains of SHP2 showed strong and preferential binding to the phosphorylated PD1 ITIM and ITSM motifs over unphosphorylated ITIM/ITSM with the N terminal SH2 domain showing strong binding to both motifs by ELISA whilst the C-terminal SH2 domain demonstrated a mild preference for the ITSM. The dual SH2 domains of SHP1 also displayed binding to both the PD1 phospho-ITIM and phospho-ITSM although with a lower EC50 for both (Table 1). The C- terminal SH2 domain showed similar binding to the phospho-ITSM sequence as that of SHP2 whilst the N-terminal domain appeared unable to interact with either motif (Figure 1B).

**Table 1:** EC50 of SH2 domains of SHP1 and SHP2 binding to phospho-ITIM and phospho-ITSM motifs of PD1. ND: not detectable.

| EC<sub>50</sub> (nM) | pITIM | pITSM |
| --- | --- | --- |
| SHP1 - C-Term | 33.72 | 1.689 |
| SHP1 - N-Term | ND | 266.3 |
| SHP1 - Dual | 21.39 | 15.27 |
| SHP2 - C-Term | 5.556 | 1.852 |
| SHP2 - N-Term | 12.04 | 4.583 |
| SHP2 - Dual | 1.686 | 1.238 |

This pattern of binding was only partially recapitulated by surface plasmon resonance with the SH2 domains of SHP1 showing little to no interaction with the phospho-ITIM in any format (Figure 1C). Notably, using this method, we were also unable to detect binding of the SHP2 C-terminal SH2 to the phospho-ITIM. In accordance with the results from the ELISA, the N-terminal domain of SHP1 showed little binding to either phosphopeptide and binding was not observed for any SH2 domain construct to the unphosphorylated ITIM or ITSM (Supplementary Figure 1).

### Co-expression of truncated SHP2 blocks PDL1/PD1 mediated CAR T inhibition

Dual SH2 domains from SHP1 or SHP2 (dSHP1, dSHP2) were then co-expressed with a second generation CD28-CD3ζ anti-GD2 CAR^12^ and the marker gene RQR8^13^ separated by viral 2A peptide sequences. To simulate exhaustion, in some conditions, T cells were additionally transduced to express PD1 (Figure 2A). T cells were challenged with SupT1 cells (which lack GD2), SupT1 cells engineered to express GD2 and SupT1 cells engineered to express both GD2 and PDL1 (Supplementary Figure 2).

**Figure 2:**
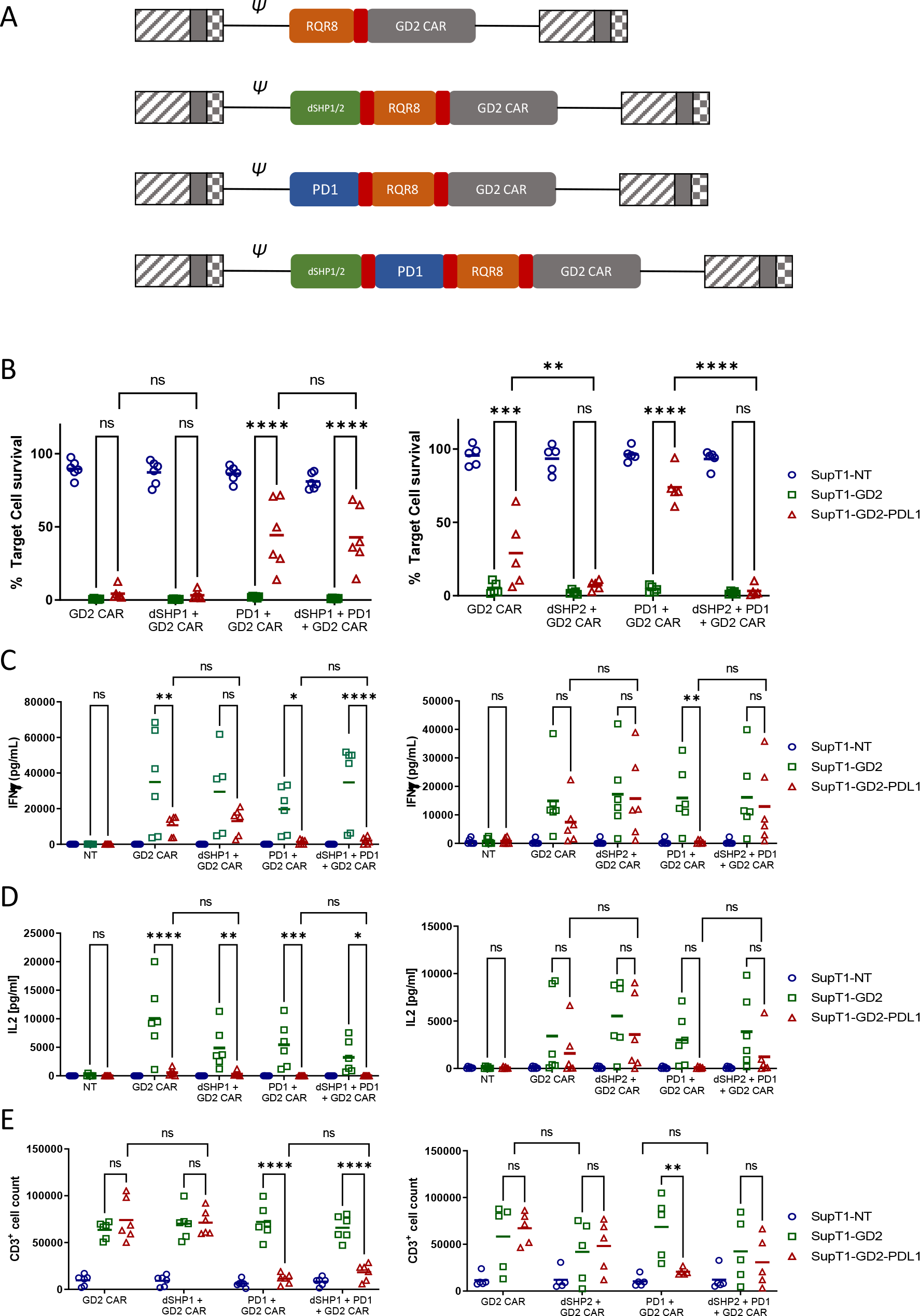
**dSHP2 but not dSHP1 alleviates PD1-mediated suppression of CAR T activity. A**) Schematic of CAR constructs. Retroviral constructs were designed consisting of an anti-GD2 CAR with a CD28-CD3ζ endodomain linked to RQR8 as a marker of transduction and including PD1 and/or dSHP1 or dSHP2 separated by viral 2A sequences (shown in red). (**B**) Effect of dSHP1 (left panel) and dSHP2 (right panel) on PD1/PDL1-mediated suppression of CAR cytotoxicity. CAR T cells were co- cultured with 2x10^5^ SupT1 cells lacking antigen expression (SupT1-NT) or engineered to express GD2 alone (SupT1-GD2) or in the presence of PDL1 (SupT1-GD2-PDL1) at an effector target ratio of 1:4 for 72 hours. Surviving target cells were enumerated and normalized to the respective co-cultures with non-transduced T cells (100%). **C-D**) Effect of dSHP1 (left panel) and dSHP2 (right panel) on PD1/PDL1-mediated suppression of IFNγ and IL2 secretion. Supernatants from (B) were harvested and cytokine secretion measure by ELISA. **E**) Effect of dSHP1 (left panel) and dSHP2 (right panel) on PD1/PDL1-mediated suppression of proliferation. CAR T cells were labelled with Cell Trace violet and incubated with 2x10^5^ of the indicated SupT1 target cells at an E:T ratio of 1:2 for 4 days and CAR T cells enumerated. Bars represent the mean of 5-6 biologically independent replicates. Statistical significance was measured by two-way ANOVA.

PDL1 expressing targets were less efficiently killed by CAR T cells than those expressing GD2 alone, however the immunosuppressive effect of target cell PDL1 expression was overcome by dSHP2, with complete restoration of killing. dSHP1, in contrast, had no effect (Figure 2B). The cytotoxic capacity of the CAR alone appeared to diminish upon serial exposure to target cells (Supplementary Figure 3A-B), however this capacity was maintained when dSHP2 was present suggesting that target cell-expressed PDL1 may be responsible. The effect was more pronounced when exogenous PDL1 was expressed on the target cells and the expression of PD1 alongside the CAR further impaired the CAR-mediated cytotoxicity.

We also examined the secretion of cytokines from CAR T cells in these co-cultures. IFNγ secretion was inhibited by PDL1 even in the absence of exogenous PD1 however this effect was enhanced when CAR T cells co-expressed PD1 (Figure 2C). Inhibition of IFNγ was ameliorated by co- expression of dSHP2 in challenge against SupT1-GD2-PDL1 target cells, but not by co-expression of dSHP1. A similar pattern was observed for IL2. However, in the case of both cytokines, restoration of secretion by dSHP2 was incomplete: failing to reach the levels measured in the absence of target cell PDL1. Proliferation of CAR T cells was suppressed by PDL1 on target cells in the presence of exogenous PD1 on the T cells and restoration with SHP2 was limited. Additionally, proliferation was diminished for cells expressing dSHP2 alongside the GD2 CAR upon serial rechallenge with antigen-expressing target cells, however we could detect no difference in the degree of apoptosis CAR T cells undergo following exposure to target cells regardless of the presence or absence of dSHP2 (Supplementary Figure 3C-D).

Subsequently, we investigated the impact of genomic ablation of SHP2 on PDL1 mediated inhibition in comparison to the dSHP2 module. Nucleofection of Cas9 RNP complexes containing SHP2- directed sgRNA into CAR T cells prior to transduction, resulted in greater than 80% indel formation and a significant decrease in SHP2 protein detected by Western blot after 6 days (Supplementary Figure 4A-C). As before, PDL1 expressed on target cells inhibited the T cells expressing the GD2 CAR alongside PD1 and this inhibition was relieved by dSHP2 (Supplementary Figure 4D).

However, this relief of inhibition was substantially less for CAR T cells treated with non-targeted or SHP2-targeted Cas9 RNPs.

We next sought to compare dSHP2 with Pembrolizumab by carrying out co-culture assays of T cells expressing GD2 CAR with either dSHP2 co-expression or in the presence of increasing concentrations of Pembrolizumab (Supplementary Figure 5). Restoration of killing, IFNγ release and proliferation with co-expression of dSHP2 was comparable to that seen with saturating amounts of Pembrolizumab.

### Co-expression of truncated SHP2 is applicable to different CAR architectures and CARs recognizing different targets

We also sought to determine whether these effects could be replicated for a CAR with a 41BBz endodomain. Above experiments were repeated with a 41BB-CD3ζ anti-GD2 CAR (Supplementary Figure 6). As before we observed inhibition of killing and cytokine release in response to targets expressing PDL1, with this effect being more pronounced with T cells having exogenous PD1 expression. Once again dSHP2 could provide a restoration of function, however this restoration was incomplete for both cytotoxicity and cytokine secretion perhaps indicating a greater sensitivity of this 41BBz CAR to PD1-mediated inhibition.

We next tested dSHP2 in the context of a CD19 CAR^14^ with a CD28-CD3ζ endodomain. We performed co-culture experiments with SupT1 cells engineered to express CD19, or CD19 and PDL1 and in some conditions PD1 was co-expressed in CAR T cells (Supplementary Figure 7A). Targets expressing only CD19 were efficiently killed by CD19 CAR T cells and triggered IFNγ release and CAR T proliferation (Supplementary Figure 7B-D). However, against targets which co-expressed PDL1, and in the context of exogenous PD1 expression, CAR T mediated killing, cytokine secretion and proliferation were all significantly reduced. The expression of dSHP2 within the CAR T cells fully restored cytotoxicity and proliferation, and partially restored cytokine release (Supplementary Figure 7B-D).

### Truncated SHP2 and CAR co-expression does not result in exhaustion or increased differentiation

We sought to next determine the effect of dSHP2 expression on T cell activation, including exhaustion and differentiation, utilizing increasing concentrations of plate-bound anti-CAR-idiotype antibody for activation. Following 24 hours upregulation of CD69, CD25 and PD1 were comparable between cells expressing CAR alone and CAR with dSHP2 (Supplementary Figure 8A-C). Additionally, no differences were observed in the memory or exhaustion profile of the cells following 72 hours of stimulation (Supplementary Figure 8D-E).

Transcriptional analysis was also performed. In T-cells expressing the CAR alone or the CAR co-expressed with dSHP2, clustering was seen based on the activation status with activated and non-activated cells forming defined and separate clusters for each construct (Supplementary Figure 9A-B left panels). However, when we compared activated cells expressing the CAR alone of the CAR with dSHP2 we found that the clusters identified corresponded to each donor and reflected donor variation rather than variation between the CAR constructs (Supplementary Figure 9A-B right panels). This indicated that variation between donors was more significant than differences between constructs. A volcano plot of differentially expressed genes indicated that no individual gene reached the threshold for a significant difference in expression in activated cells expressing or lacking the dSHP2 model indicating that dSHP2 did not exert a significant effect on gene expression following CAR T activation (Supplementary Figure 9C).

### Truncated SHP2 and other inhibitory receptors

In order to understand the potential interaction of SHP2 with other ITIM/ITSM receptors, we once again used ELISA to measure the interaction of the tandem SH2 domains of SHP2 (dSHP2) to phosphorylated a peptides from inhibitory receptor endodomains and their non-phosphorylated variants (Supplementary table 1). We tested the interaction of SHP2 with phosphorylated and non-phosphorylated ITIM or ITSM motifs derived from PAG, TIGIT, BTLA, 2B4 and CTLA-4 (Figure 3A-E). In the case of TIGIT peptide 1 and CTLA4 peptide 19, two potential tyrosine phosphorylation sites were present and binding was tested to peptide having both site phosphorylated or each one individually. We observed promiscuous binding of dSHP2 to phospho-peptides from all of the receptors tested with the exception of PAG.

**Figure 3:**
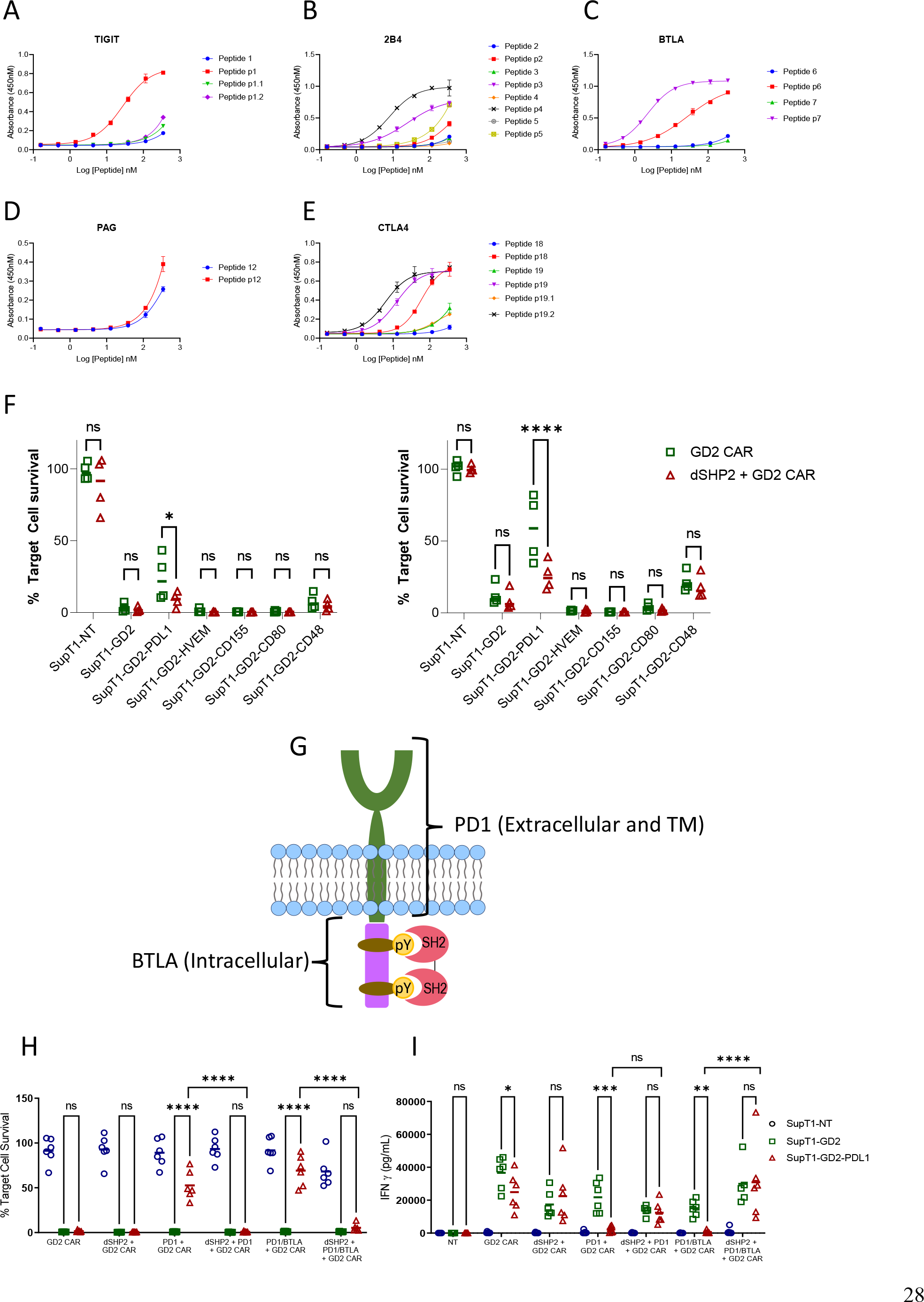
**Interaction of dSHP2 with immunoinhibitory T cell receptors. A-E**) Binding of dSHP2 to phosphorylated peptides derived from the endodomains of immunoinhibitory T cell receptors. Plates were coated with phosphorylated or non-phosphorylated peptide and the indicated concentration and the purified, His-tagged SH2 domains of dSHP2 were allowed to bind overnight. Binding was measured through detection of the His tag. F) Effect of inhibitory ligand expression on target cells on CAR T cytotoxicity. CAR T cells expressing the GD2 CAR alone or alongside dSHP2 were co-cultured with 5x10^5^ SupT1 target cells expressing no antigen (SupT1-NT) or engineered to express GD2 alone or in combination with the indicated inhibitory ligand at an E:T ratio of 1:4 (left panel) or 1:8 (right panel). Live target cells were enumerated after 6 days and normalized to the respective co-culture of the target cells with non-transduced T cells (100%). G) Schematic of the PD1/BTLA chimeric constructs. A chimeric construct was designed consisting of the ecto- and transmembrane domain of PD1 linked to the intracellular domain of BTLA. H-I) Effect of dSHP2 on immune suppression mediated by PD1/BTLA chimera. CAR T cells expressing the GD2 CAR in the presence or absence of dSHP2 and full length PD1 or the PD1/BTLA chimera were co-cultured with SupT1 target cells expressing no antigen (SupT1-NT) or engineered to express GD2 alone or in combination with PDL1 at an E:T ratio of 1:4 for 72 hour. Live target cells were enumerated and normalized to the respective co-culture of the target cells with non-transduced T cells (100%). Supernatants were harvested from these co-cultures and levels of IFNγ measured by ELISA (I). Bars represent the mean of 4-6 biologically independent replicates. Statistical significance was measured by two-way ANOVA.

In order to investigate the effect of dSHP2 on inhibition mediated by receptors other than PD1 we transduced GD2-SupT1 with a range of inhibitory ligands which interact with the inhibitory receptors BTLA (HVEM), TIGIT (CD155), CTLA4 (CD80) and 2B4 (CD48) and mediate inhibition of T cell function. We then carried out co-cultures to measure the effect of inhibitory ligand expression on cytotoxicity mediated by the GD2 CAR in the presence or absence of dSHP2. Surprisingly, we were unable to observe any inhibition meditated by target cell ligand expression other than with PDL1. In fact, we observed a mild enhancement of CAR T cytotoxicity induced by HVEM, CD155 and CD80 suggesting a stimulatory effect on the T cells in this co-culture.

We next sought to determine whether inhibitory effects were being masked by the presence of stimulatory receptors which could also bind target cell expressed ligands. Since BTLA is well characterized and its endodomain showed strong binding to dSHP2, we generated a chimeric construct consisting of the PD1 ecto- and transmembrane domains linked to the endodomain of BTLA (Figure 3G). We co-expressed this construct in T cells alongside the GD2 CAR in the presence or absence of dSHP2. Co-cultures of transduced CAR T cells were performed with target cells expressing GD2 in the presence or absence of PDL1 (Figure 3H-I). As a control for inhibition, we utilized the CAR co-expressed with PD1 as described previously. In this format, we observed inhibition of CAR T cell killing and cytokine release by PD1/BTLA expressing T cells in response to targets expressing PDL1 and this inhibition was countered by dSHP2.

### Truncated SHP2 enhances CAR T cell activity in an in vivo model of PDL1 expressing tumor

Next, we tested the efficacy of dSHP2 in an in vivo model utilizing a CD19-expressing Nalm6 model and the CD19 CAR. In order to measure the effect of PDL1 on CAR efficacy in vivo we engrafted either wild type Nalm6 cells or Nalm6 cells engineered to express PDL1 in NSG mice. and treated with mock transduced T cells (NT), T cells expressing the CD19 CAR alone or in combination with dSHP2 (Figure 4 and Supplementary Figure 10). We observed that T cells expressing the CAR alone and the CAR + dSHP2 could prevent tumor growth (Figure 4A) however by approximately day 20 tumors began to regrow in the cohort treated with CAR + dSHP2 and several mice had to be culled (Figure 4C). The presence of PDL1 on the Nalm6 cells appeared to greatly reduce the efficacy of T cells expressing CAR alone to mediate tumor rejection and all mice in this cohort failed to survive past D20 post CAR T injection (Figure 4B). However, 3/6 mice treated with CAR + dSHP2 displayed delayed tumor growth followed by rejection in the later stages of the model and were still alive at day 34 (Figure 4D) indicating the dSHP2 was able to counter the suppressive effect of PDL1 and enhance mouse survival and CAR T function in this context. Interestingly enumeration of CAR T cell numbers suggested that the presence of dSHP2 resulted in a reduction of CAR T expansion which may explain the incomplete survival of the CAR and dSHP2 cohort engrafted with wild type Nalm6 cells (Figure 4E-F).

**Figure 4:**
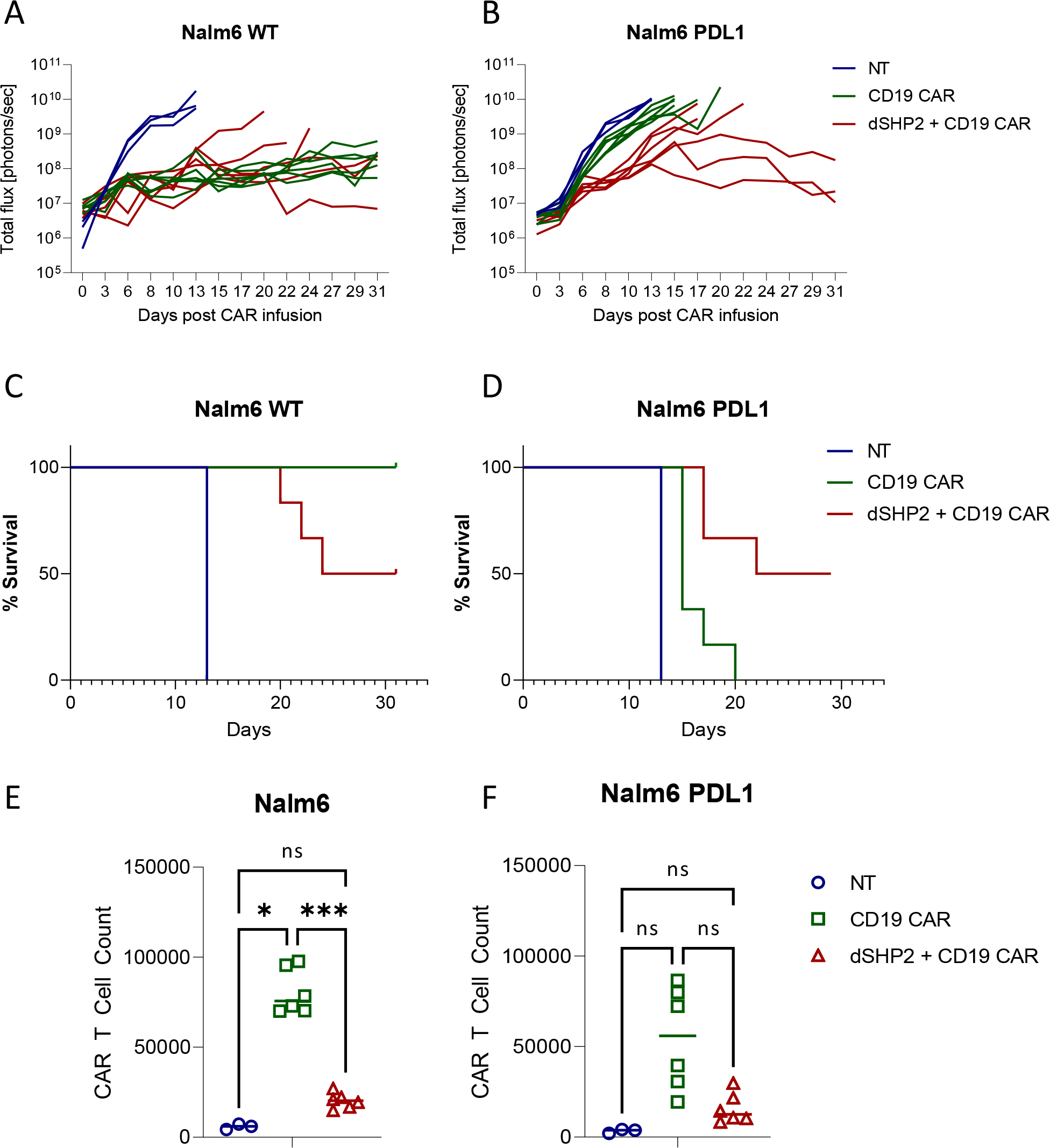
In vivo effect of dSHP2 co-expressed with a CD19 CAR. NSG mice were engrafted with 0.5x10^6^ Nalm6 cells engineered to express firefly luciferase and where indicated PDL1. Four days later mice were injected with 1x10^6^ CAR T cells expressing the CD19 CAR or the CD19 CAR alongside dSHP2 or an equivalent amount of non-transduced NT T cells. **A-B**) Growth of tumor was monitored by bioluminescent imaging. Each line represents an individual animal. **C-D**) Kaplan-Meier curve of survival. **E-F**) CAR T cells were enumerated in the bone marrow through flow cytometry detecting CAR expression through staining with an anti-idiotype antibody specific for the binding domain of the CAR.

In light of the observed reduction in proliferation of CAR T cells expressing dSHP2 we examined the effect of this construct on pathways involved in cytokine signaling. Exposure to IL2 promoted ERK phosphorylation in NT T cells and T cells expressing a CD19 CAR alone however this was substantially reduced in the presence of dSHP (Supplementary Figure 11A and B). Additionally, activation of CAR T cells with anti-CAR idiotype antibody combined with IL2 lead to the phosphorylation of ribosomal protein S6 and this was also reduced when dSHP was present (Supplementary Figure 11C and D).

## Discussion

SHP-1 and SHP-2 are members of the Src-homology 2 domain (SH2)-containing PTP family unique in that they possess protein tyrosine phosphatase activity^15^. Within immune cells, SHP-1 and SHP-2 are recruited to the cell membrane upon phosphorylation of inhibitory immune receptor endodomains^16, 17^ where they dephosphorylate, and hence inactivate, key mediators of immune cell activation leading to immunosuppression^18, 19^. Consequently, they represent a key point of convergence and pinch-point for the transmission of immune inhibitory signals.

SHP1 and SHP2 display a high sequence identity and are structurally similar composed of two SH2 domains, a catalytic protein tyrosine phosphatase (PTP) domain and a C-terminal tail. The N- terminal SH2 domain regulates SHP activity by obstructing the catalytic site of the PTP domain^20^. Association of the N-terminal SH2 domains with phosphorylated ITIM/ITSM motifs in the cytoplasmic tail of cell surface inhibitory receptors relieves this autoinhibition and localizes SHP1/2 proximal to the membrane within an immune synapse allowing them to dephosphorylate CD28, Lck and CD3ζ^19^. We hypothesized that expression of truncated forms of these proteins consisting only of the tandem SH2 domains and lacking the inhibitory phosphatase domain would inhibit SHP1/2 signaling.

In our initial experiments we explored the interaction of the SH2 domains of SHP1 and SHP2 with the ITIM and ITSM motif of PD1 using surface plasmon resonance and ELISA. We found that the tandem SH2 domains of SHP2 and the individual C- and N-terminal domains bind to both the ITIM and ITSM motifs of PD1. In contrast, in our hands, the N-terminal domain of SHP1 showed no binding to either sequence suggesting that the binding observed for the dual SHP1 SH2 domains is mediated solely by the C-terminal domain. However, the N-terminal SH2 domain may not fold correctly when expressed in isolation resulting in an apparent lack of binding, thus, we cannot rule out that the N-terminal domain might contribute to binding of PD1 ITIM/ITSM when expressed in tandem with the C-terminal domain. These data are somewhat consistent with that of others: Yokosuka et al., demonstrated that SHP2 but not SHP1 co-localized with PD1, whilst the binding of SHP2 was more dependent on the ITSM than the ITIM^21^. Using surface plasmon resonance, Xu et al., demonstrated that the N-terminal SH2 domain of SHP2 shows a moderate preference for the ITSM over the ITIM whilst the C-terminal domain strongly prefers the ITSM^22^. Additionally, Xu showed that the C-terminal SH2 domain of SHP1 bound more strongly to the ITSM with little binding to the ITIM. In contrast to our data however they observed binding of the N-terminal domain of SHP1 to both the ITIM and ITSM.

In order to explore whether truncated variants of SHP1 and SHP2 could prevent PD1-mediated inhibition of CAR T cells, we co-expressed them with a GD2-specific CAR. CAR T cells were challenged with target cells expressing GD2, in the absence or presence of PDL1. We found that expression of PDL1 on the target cells inhibited CAR T function and this effect was increased in the presence of exogenously expressed PD1. In these co-cultures, cytotoxicity, secretion of IFNγ and IL2 and proliferation were all reduced by the presence of target cell-expressed PDL1. Co-expression of dSHP2 was capable of reversing this suppression to varying degrees: cytotoxicity and IFNγ secretion were restored to levels observed against target cells lacking PDL1 whilst the effects on IL2 secretion and proliferation were more modest. The partial effect on IL2 secretion may be due to more stringent requirements for IL2 secretion which has been shown to be dependent on prolonged TCR signaling^23^. By contrast dSHP1 conveyed no restoration of any CAR T function. While initial experiments utilized a CD28-ζ CAR, a similar effect of dSHP2 co-expression was also observed using a GD2 CAR with a 41BB-ζ endodomain. Additionally, similar effects were observed when dSHP2 was co- expressed with a CD19 CAR.

Expression of dSHP2 appeared to confer a greater relief of PDL1-mediated inhibition of CAR T cell function than CRISPR/Cas9-mediated knock out of the SHP2 gene possibly due to redundant mechanisms which may exist in the cells which are capable of compensating for SHP2 loss but unable to compete with the interaction of dSHP2 with PD1. We also compared the effect of dSHP2 to blocking PD1 using the antibody pembrolizumab and found comparable effects in terms of restoring cytotoxicity and proliferation and a slightly greater restoration of IFNγ secretion at the highest concentration of blocking antibody.

The inability of dSHP1 to relieve PD1-mediated inhibition was consistent with emerging understanding of SHP1 and 2 binding to ITIM and ITSM motifs within inhibitory receptor endodomains. SHP2 but not SHP1 has been shown to associate with PD1 and to accumulate in PD1 microclusters upon ligation by PDL1^21^. Furthermore, PD1-PDL1-mediated suppression of IL2 secretion in jurkat cells was unaffected by SHP1 knockout but was partially alleviated by knockout of SHP2^22^. Whilst we and others have shown that the tandem SH2 domains of SHP1 can interact with the ITSM and to a weaker degree the ITIM of PD1^17, 19^ this interaction is of a lower affinity than that displayed by the tandem SH2 domains of SHP2, particularly towards the ITSM motif. This suggests that dSHP1 lacks the affinity necessary to out-compete endogenous full length SHP2 and relieve PD1-mediated inhibition. The lower affinity of SHP1 for PD1 can be explained in part by its structure: modelling suggests that SHP1 possesses a hydrophobic pocket within the N-terminal SH2 domain which serves to stabilize the interaction between SHP1 and ITIM motifs providing these motifs possess a medium-sized non-polar residue at position +1 respective to the phospho-tyrosine^22^. Such a residue is not present within the PD1 ITIM resulting in a loss of this stabilizing interaction and a lower affinity of PD1 for SHP1.

In the immune system, both SHP1 and SHP2 mediate signaling from inhibitory receptors^17, 21^. In addition to the role SHP2 plays in the transmission of inhibitory signals and T cell exhaustion in response to chronic stimulation, it may also be involved in signaling pathways downstream of cytokine receptors^24^. As such the knock-out or inhibition of these proteins may have varying effects on T cells. We hence investigated broader consequences of expression of dSHP2 on T cell function. We tested activation and phenotype across a range of increasing concentrations of immobilized CAR- activating antibody. We observed no effect on the upregulation of CD69, CD25 or PD1 albeit with a non-significant trend to a reduction in the levels of the latter. Additionally, we saw no significant differences in transcriptomics profile of T cells expressing the CAR alone or the CAR alongside dSHP2 following activation through the CAR.

Modulating SHP2 in T cells has been previously explored: knockout of the SHP2 gene using Cre- recombinase under the control of the Lck promoter or expression of a phosphatase-null mutant in CD2-expressing T cells both resulted in reduced proliferation of mature T cells upon stimulation^25, 26^; however, a subsequent study using Cre-recombinase induced by the CD4 promoter showed that SHP2-deficient T cells had a greater proliferative capacity than wild-type T cells^27^. These studies found that secretion of cytokines in SHP2-KO T cells was impaired compared to wild type cells. In contrast, an earlier study expressing the phosphatase-null dSHP2 under the control of the CD2 promoter found that proliferation and IL2 release were unaffected following stimulation through the TCR and that peripheral blood T cells expressing the dominant negative SHP2 had higher levels of activation markers^28^. Equally suppressing SHP2 activity with the allosteric inhibitor SHP099 resulted in an increased central memory population and increased expression of granzyme B, perforin, IFNγ, TNFα and FasL following activation whilst having no effect on proliferation of viability^29^. The differing results may be consequent to the point in T cell development at which SHP2 activity is inhibited or the strength and duration of the inhibition.

Several inhibitory receptors in addition to PD1 may signal through SHP1/2. SHP1 and SHP2 have been shown display preferences in terms of the inhibitory receptors to which they respectively bind^16, 17^ with isolated SH2 domains showing greater promiscuity^19, 21, 22^. In agreement with this we demonstrated that the individual and tandem SH2 domains of SHP2 appear to interact with the phosphorylated ITIM and ITSM motifs from several inhibitory receptors with varying affinities.

Based on this we investigated whether we could detect inhibition of CAR T activity mediated by these receptors and their respective ligands and whether any inhibition could be relieved by dSHP2. Interestingly none of the inhibitory ligands tested displayed inhibitory activity towards the CAR T cells when expressed on tumor cells and tested in co-cultures at different effector-target ratios. Both HVEM and CD155, the ligands for BTLA and TIGIT respectively, appear to enhance cytotoxicity. However, when we tested a chimeric version of BTLA, which could be triggered by PDL1 rather than HVEM, we observed inhibition of the CAR T cells under conditions in which endogenous PD1 appeared to have no effect and this inhibition was relieved by dSHP2. Although in our work we did not detect an effect of dSHP1 on CAR T cell inhibition mediated by PD1 the specificity displayed by SHP1 and SHP2 for different inhibitory receptors suggests that dSHP1 may relieve immunosuppression caused by receptors other than PD1.

We tested the effect of dSHP2 expression in CD19 CAR T cells with an in vivo model utilizing Nalm6 cells, which express CD19. Expression of dSHP2 reduced the capacity of CAR T cells to reject tumor in the absence of PDL1 expression with several relapses in the respective cohort and a concomitant reduction in survival over the course of the model. In contrast, when Nalm6 cells engineered to express PDL1 were used, dSHP2 co-expression enhanced CAR T efficacy. Consistent with studies suggesting that SHP2 activity supports T cell proliferation^25, 26^ dSHP2 expression reduced the numbers of CAR T cells irrespective of the presence or absence of PDL1 on the tumor. The failure to achieve full rejection of the tumor in the cohort expressing PDL1 may be a consequence of reduced proliferation or indicative of redundant but inefficient mechanisms mediating inhibition of CAR T activity through PD1 in the absence of SHP2. However constitutive blockade of PD1 signaling may lead to an impairment of anti-tumor efficacy and in models of chronic LCMV infection genetic deletion of PD1 led to an accumulation of terminally differentiated T cells^33, 34^. In addition, transient rest of CAR T cells through a cessation of CAR signaling has been shown to be beneficial by enhancing the efficacy of CAR T cell therapy^35^. Additionally, we noted that dSHP2 appeared to dampen pathways involved in T cell activation following exposure of the T cell to IL2 alone or combined with stimulation through the CAR potentially explaining the reduced proliferation observed for CAR T cells expressing dSHP2 in vivo and upon serial rechallenge with target cells in vitro. Hence constitutive expression of dSHP2 may lead to a reduction in overall responses which is consistent with our in vivo observations.

In a therapeutic setting, relief of T cell inhibition is most commonly achieved through administration of blocking antibodies specific for individual inhibitory. Engineered cell therapies afford more discrete methods of manipulating inhibitory pathways as the consequences of this manipulation are limited to within the immune cell itself. Current efforts have focused on individual surface receptors such as ablation of PD1 through the expression of siRNA or genomic editing^30, 31^, or expression of dominant negative and “switch receptor” variants of PD1^9, 32^. Many approaches aimed at preventing immunosuppression are limited in that they are only capable of targeting a specific receptor-ligand pair within an immunosuppressive microenvironment exhibiting a diverse array of inhibitory receptors. Systemic blockade of multiple inhibitory pathways may result in autoimmunity. Given that activation of T cells within the tumor microenvironment and, as a consequence their ability to mediate tumor rejection, is a balance between the collective activity of ITAMs and ITIM/ITSMs within these cells, development of approaches which can block pinch-points in signaling pathways such as SHP1 and SHP2 may result in more effective immunotherapies.

## Author Contributions

ST and MP conceived the project. JT, AB, IG, MR, MS, CW, WL, PP, SS, EK, TG, CM, MEK, and VB performed the in vitro and in vivo experiments. ST, JS and MF designed the experiments and reviewed the data. ST wrote the manuscript.

