## Supplemental Figures for "Exploration of T-cell immune responses by expression of a dominant-negative SHP1 and SHP2"

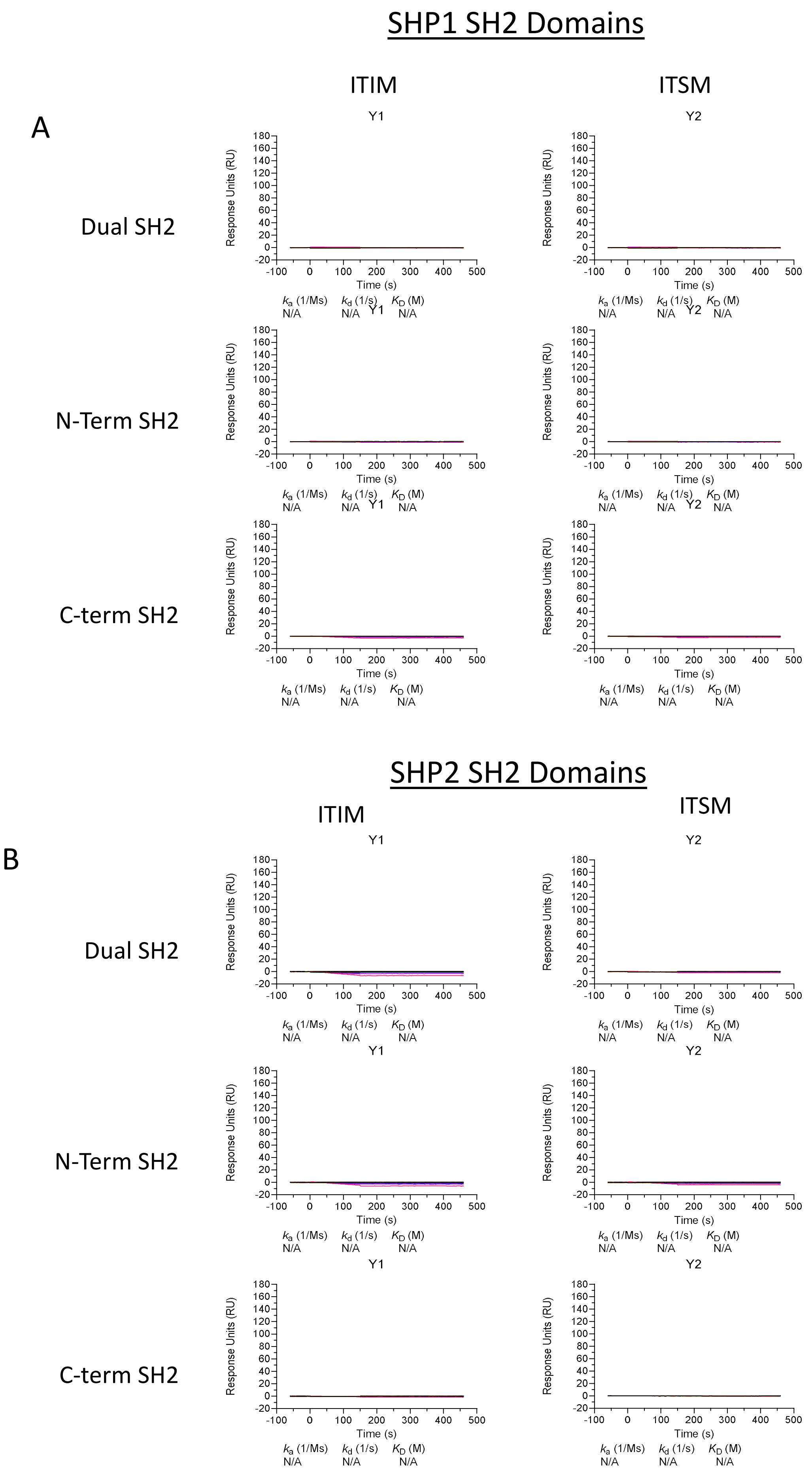


**Supplementary Figure 1: Binding of the SH2 domains of SHP1 and SHP2 to the non-phosphorylated ITIM and ITSM peptides of PD1 by Surface Plasmon resonance**

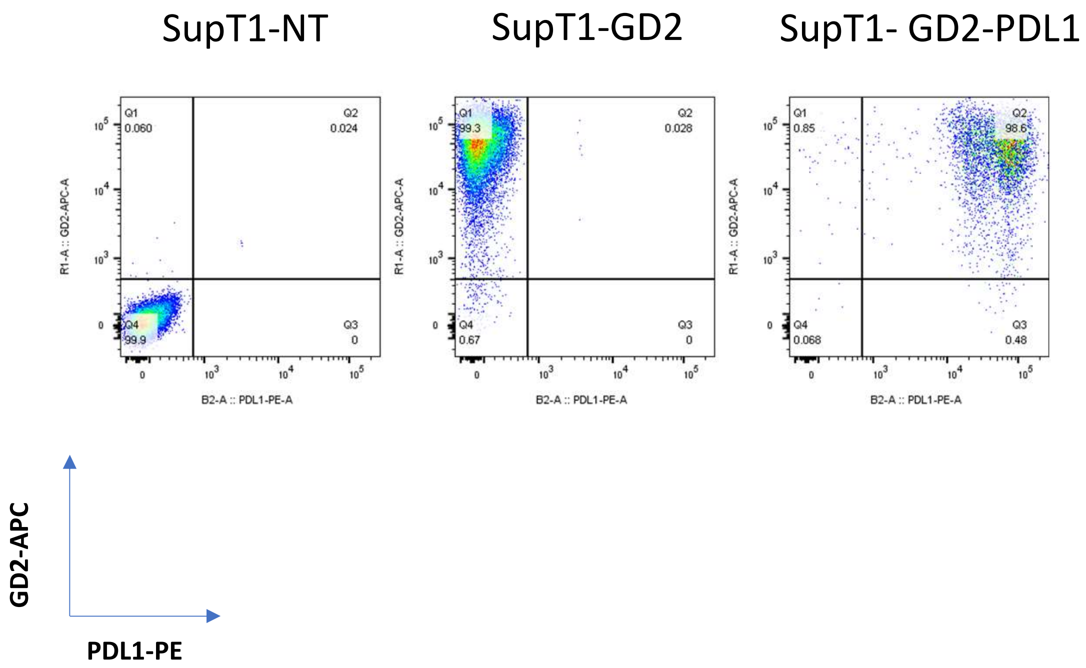


**Supplementary Figure 2: Expression of GD2 and PDL1 on engineered SupT1 target cells.**

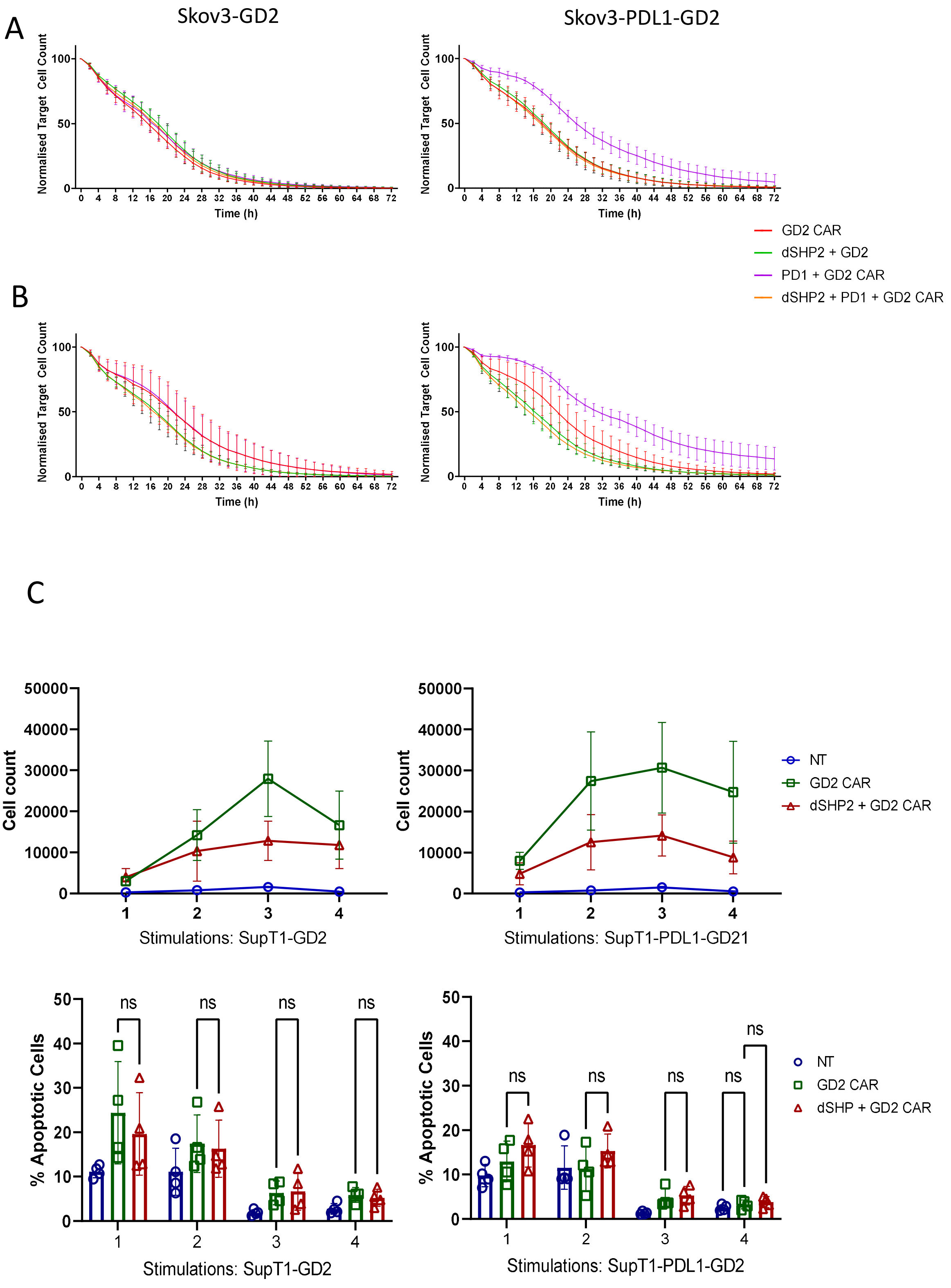


**Supplementary Figure 3: CAR T function upon serial challenge with target cells**. A-B) Serial Cytotoxicity. T cells expressing the indicated CD28-CD3ζ CAR constructs were co-cultured for 72 hours with Skov3 cells expressing GD2 (Left Panels) or GD2 and PDL1 (Right panels). T cells were then harvested and rested for 3 days before a second co-culture with Skov3 cells as before. Skov3 cell viability was monitored using an Incucyte Live Cell analysis instrument. A) First co-culture. B) Second co-culture. C) CAR T proliferation. D) Apoptosis. T cells transduced to express the GD2 CAR alone or in the presence of dSHP or left non-transduced (NT) were stimulated every 3 days with SupT1 target cells expressing GD2 alone or in combination with PDL1 and T cells enumerated and assessed for activation induced cell death via Annexin V and 7-AAD staining following each stimulation.

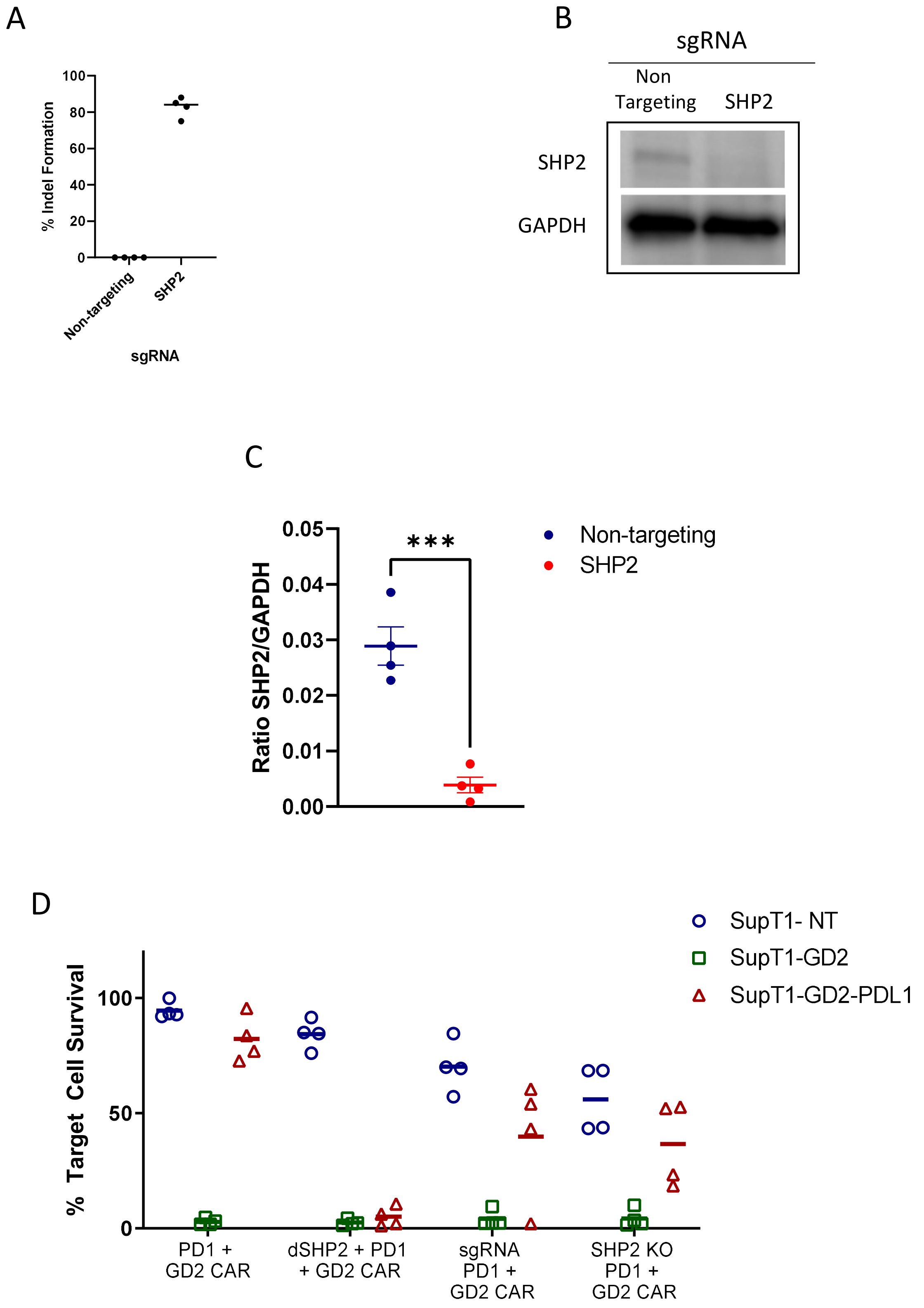


**Supplementary Figure 4**: **Comparison of dSHP2 expression with genomic excision of SHP2.** SHP2 was knocked out from T cells using CRISPR-Cas9 and cells were subsequently transduced to express the CD28-CD3ζ GD2 CAR in combination with PD1 and in the absence or presence of dSHP2 and assayed for function after 6 days against target cells expressing no antigen (SupT1-NT), GD2, (SupT1-GD2) or GD2 and PDL1 (SupT1-GD2-PDL1). A) Percentage of SHP2 knock out in four independent donors following transfection with non-targeting sgRNA or SHP2-directed sgRNA. B) Western blot showing loss of SHP2 expression following CRISPR treatment of T cells in a single donor. C) Ratio of SHP2:GAPDH protein in 4 independent donors following knockout of SHP2. D) Cytotoxicity of T cells expressing the CAR alongside PD1 in the presence or absence of dSHP2 or SHP2 KO or non targeting sgRNA.

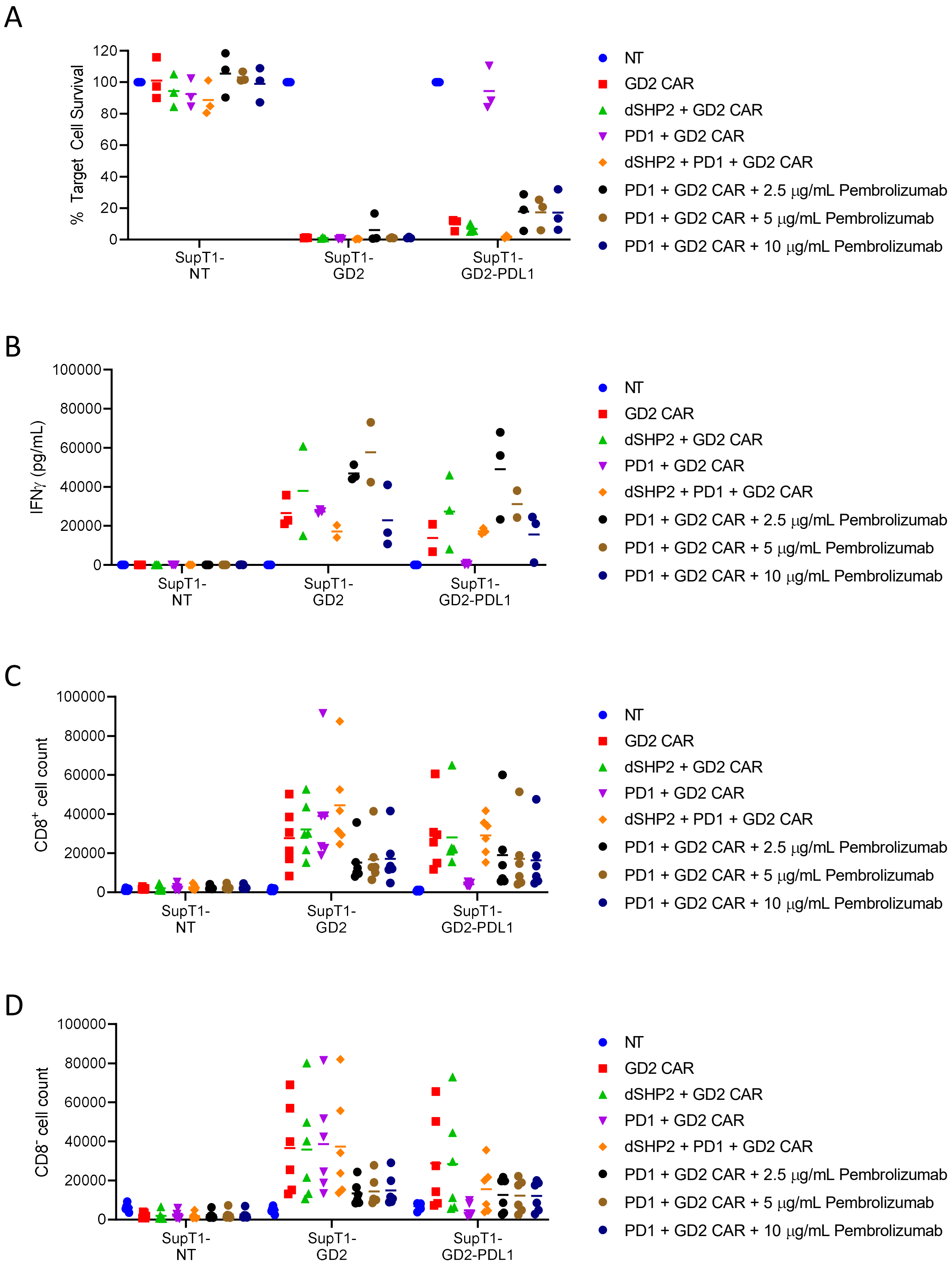


**Supplementary Figure 5: Comparison of dSHP2 with Pembrolizumab. A**) T cells expressing RQR8 and the GD2 CAR alone or in combination with PD1 and/or dSHP2 were incubated with $1x10^5$ SupT1 cells lacking antigen expression (SupT1-NT) or engineered to express GD2 alone (SupT1-GD2) or in the presence of PDL1 (SupT1-GD2-PDL1) at an effect target ratio of 1:8 for 72 hours. Prembrolizumab was included in co-cultures with T cells expression PD1 and the GD2 CAR at the indicated concentrations. Surviving target cells were enumerated and normalized to the respective co-cultures with non-transduced T cells (100%). **B**) Supernatants from co-cultures in (A) were analyzed for IFNγ by ELISA. **C**) T cells were labelled with Cell Trace Violet and co-cultured with $2x10^5$ SupT1 target cells at en E:T ratio of 1:4 for 5 days. Bars represent the mean of 3-6 biologically independent replicates

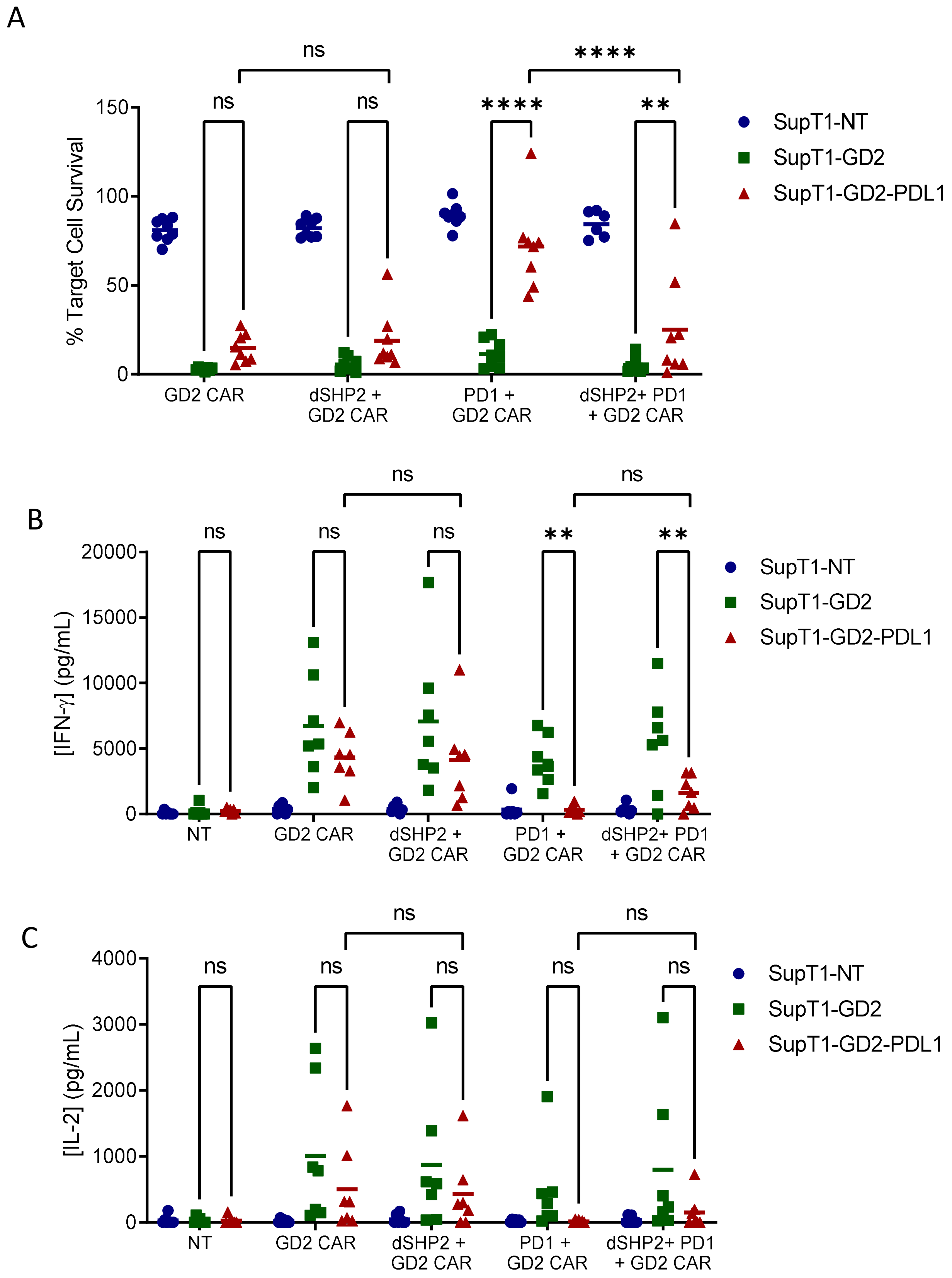


**Supplementary Figure 6: Effect of dSHP2 on alleviating immune suppression in the context of a CAR with a 41BB-CD3ζ endodomain. A**) CAR T cells were co-cultured with $2x10^5$ SupT1 cells lacking antigen expression (SupT1-NT) or engineered to express GD2 alone (SupT1-GD2) or in the presence of PDL1 (SupT1-GD2-PDL1) at an effector target ratio of 1:4 for 72 hours. Surviving target cells were enumerated and normalized to the respective co-cultures with non-transduced T cells (100%). **B-C**) Effect of dSHP1 and dSHP2 on PD1/PDL1-mediated suppression of IFNγ and IL2 secretion. Supernatants from (A) were harvested and cytokine secretion measure by ELISA. Bars represent the mean of 6 biologically independent replicates. Statistical significance was measured by two-way ANOVA.

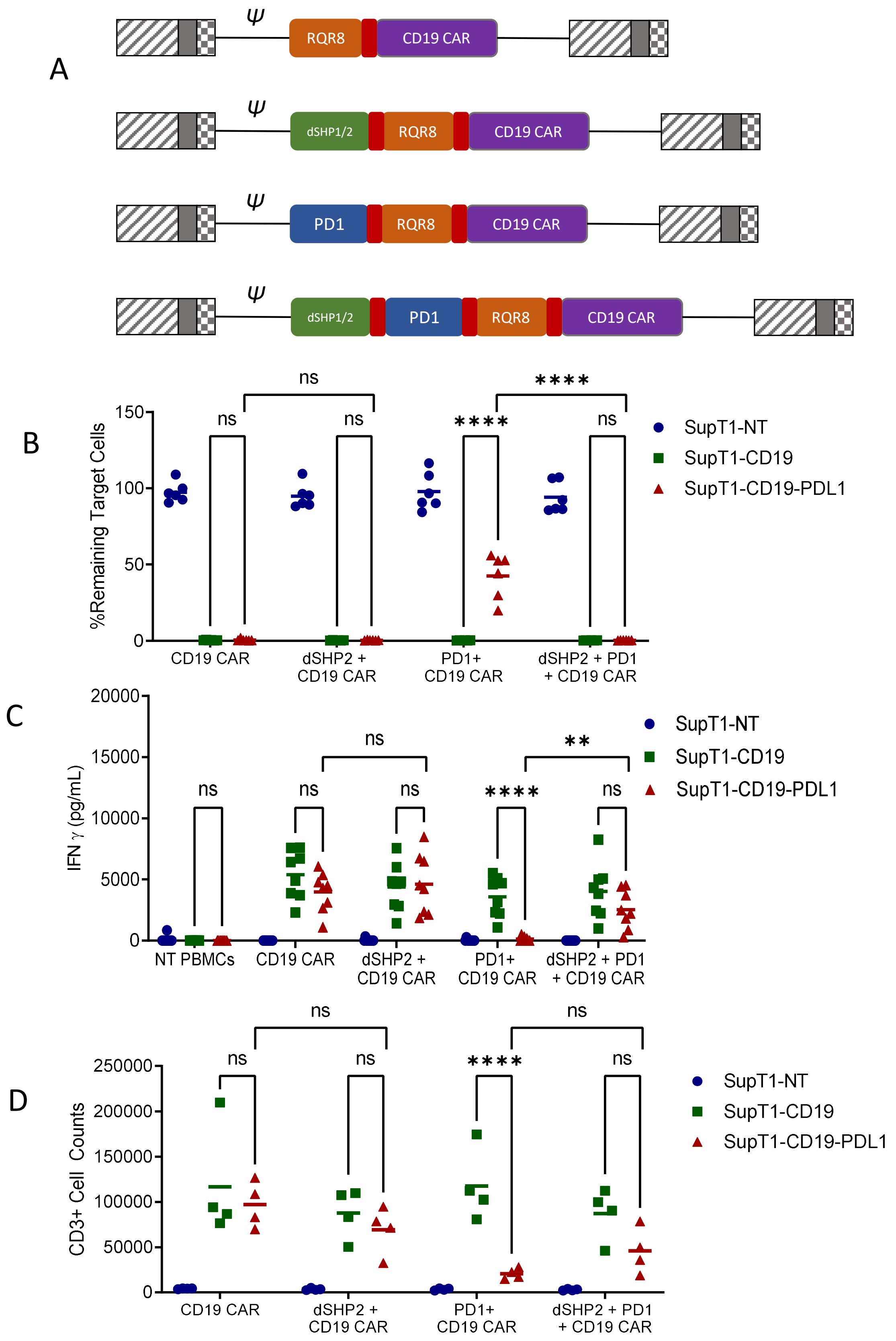


**Supplementary figure 7: Effect of dSHP2 on alleviating immune suppression in the context of a CD19 specific CAR. A**) Schematic of CAR constructs. Retroviral constructs were designed consisting of an anti-CD19 CAR with a CD28-CD3ζ endodomain linked to RQR8 as a marker of transduction and including PD1 and/or dSHP2 separated by viral 2A sequences (shown in red). (**B**) Effect of dSHP2 on PD1/PDL1-mediated suppression of CAR cytotoxicity. CAR T cells were co-cultured with $2x10^5$ SupT1 cells lacking antigen expression (SupT1-NT) or engineered to express CD19 alone (SupT1-CD19) or in the presence of PDL1 (SupT1-CD19-PDL1) at an effector target ratio of 1:8 for 72 hours. Surviving target cells were enumerated and normalized to the respective co-cultures with non-transduced T cells (100%). **C**) Effect of dSHP2 on PD1/PDL1-mediated suppression of IFNγ secretion. Supernatants from (B) were harvested and cytokine secretion measure by ELISA. D) Effect of dSHP2 on PD1/PDL1-mediated suppression of proliferation. CAR T cells were labelled with Cell Trace violet and incubated with $2x10^5$ of the indicated SupT1 target cells at an E:T ratio of 1:4 for 5 days and CAR T cells enumerated. Bars represent the mean of 4-8 biologically independent replicates. Statistical significance was measured by two-way ANOVA.

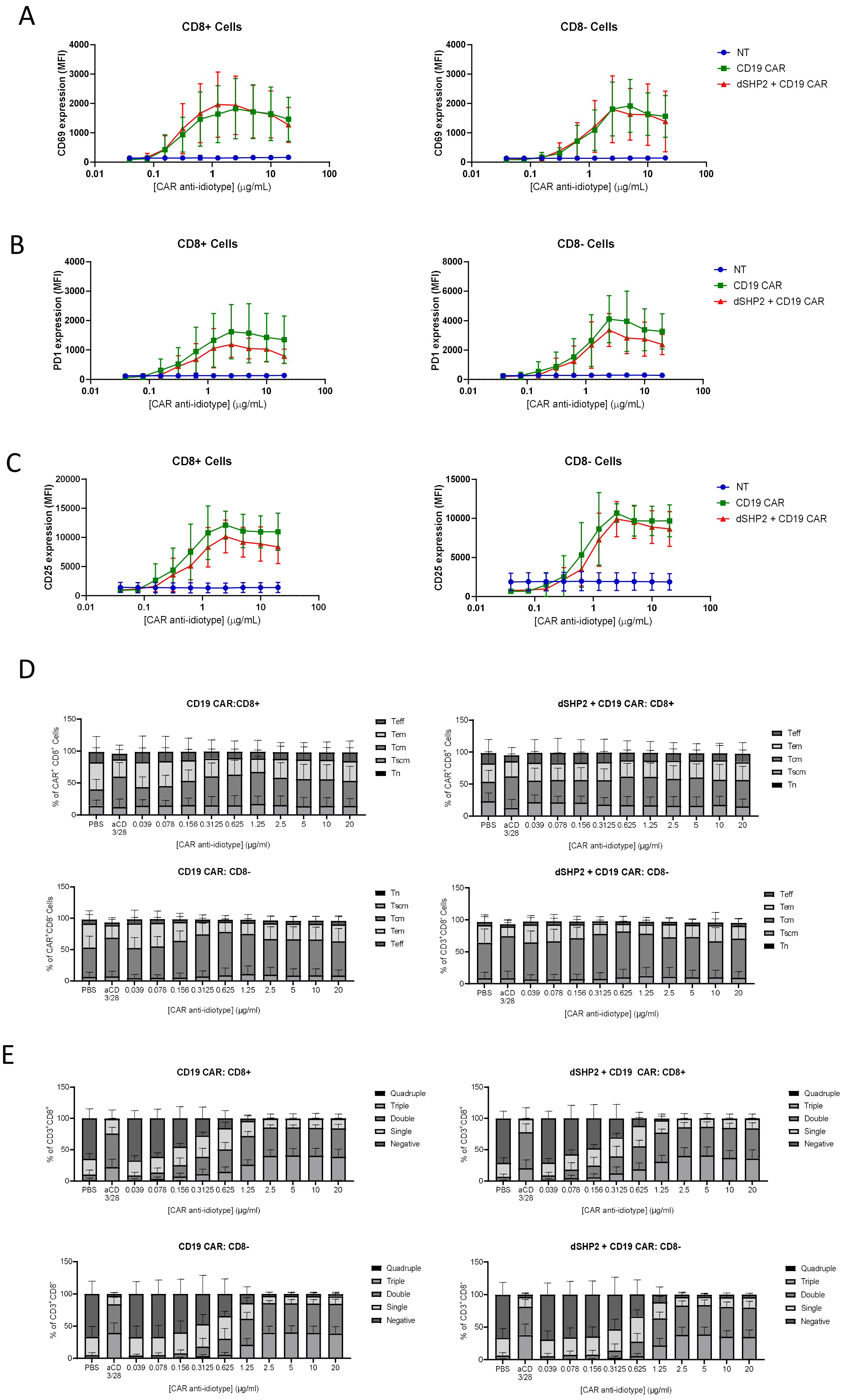
A
CD8+ Cells
CD8- Cells
CD69 expression (MFI)
[CAR anti-idiotype] (μg/mL)
NT
CD19 CAR
dSHP2 + CD19 CAR
B
CD8+ Cells
CD8- Cells
PD1 expression (MFI)
[CAR anti-idiotype] (μg/mL)
NT
CD19 CAR
dSHP2 + CD19 CAR
C
CD8+ Cells
CD8- Cells
CD25 expression (MFI)
[CAR anti-idiotype] (μg/mL)
NT
CD19 CAR
dSHP2 + CD19 CAR
D
CD19 CAR:CD8+
dSHP2 + CD19 CAR: CD8+
CD19 CAR: CD8-
dSHP2 + CD19 CAR: CD8-
Teff
Tem
Tcm
Tscm
Tn
PBS
aCD 3/28
0.039
0.078
0.156
0.3125
0.625
1.25
2.5
5
10
20
[CAR anti-idiotype] (μg/ml)
E
CD19 CAR: CD8+
dSHP2 + CD19 CAR: CD8+
CD19 CAR: CD8-
dSHP2 + CD19 CAR: CD8-
Quadruple
Triple
Double
Single
Negative
[CAR anti-idiotype] (μg/ml)

**Supplementary Figure 8: Effect of dSHP2 on CAR T cell phenotype. A-C**) Effect of dSHP2 on the upregulation of T cell activation markers. T cells expressing RQR8 and a CD19 CAR in the absence or presence of dSHP2 or non-transduced (NT) T cells were incubated on plates coated with the indicated concentration of anti-idiotype antibody specific to the CAR binding domain for 24 hours. Cells were harvested and stained for the expression of (A) CD69, (B) PD1 and (C) CD25. Graphs show the MFI of the indicated marker on CD8+ CAR (RQR8+) T cells (Left panels) or CD8- CAR (RQR8+) T cells (right panel). For NT T cells data are displayed for the total CD3+ population. Data are displayed as mean and standard deviation of 9 independent donors. **D**) Memory subtypes of stimulated CAR T cells in the absence (left panels) or presence (right panels) of dSHP. T cells were stimulated on plates coated with the indicated concentration of anti-CAR idiotype antibody for 72 hours then harvested and stained for expression of CCR7, CD45RA and CD95 on the CD8+ CAR+ cells (top panels) or CD8- CAR T cells (bottom panels). Populations are defined as Tn (CD45+/CCR7+/CD95-), Tscm (CD45+/CCR7+/CD95+), Tcm (CD45-/CCR7+/CD95+), Tem (CD45-/CCR7-/CD95+), and Teff (CD45+/CCR7-/CD95+). Cells were stimulated with anti-CD3 and anti-CD28 antibodies as a positive control. Data are shown as the mean and SD of 7 independent donors. **E**) Expression of markers of exhaustion on stimulated CAR T cells in the absence (left panels) or presence (right panels) of dSHP. T cells were stimulated on plates coated with the indicated concentration of anti-CAR idiotype antibody for 72 hours then harvested and stained for expression of KLRG1, PD1, LAG3 and Tim3 on the CD8+ CAR+ cells (top panels) or CD8- CAR T cells (bottom panels). Data are presented as the percentage of CAR T cells co-expressing 0, 1, 2, 3, or 4 exhaustion markers. Cells were stimulated with anti-CD3 and anti-CD28 antibodies as a positive control. Data are shown as the mean and SD of 7 independent donors.

### Heat Maps- Cytotoxicity

A

Stimulated vs unstimulated CD19 CAR

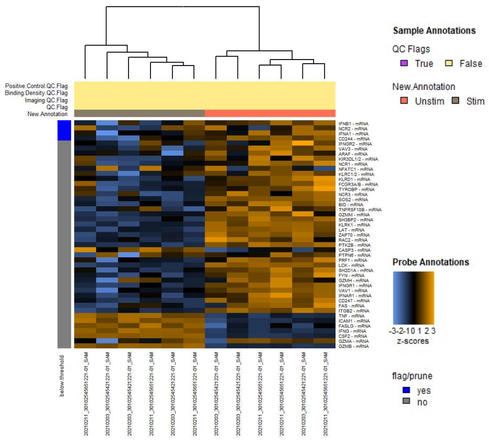


Stimulated CD19 CAR vs stimulated dSHP2 + CD19 CAR

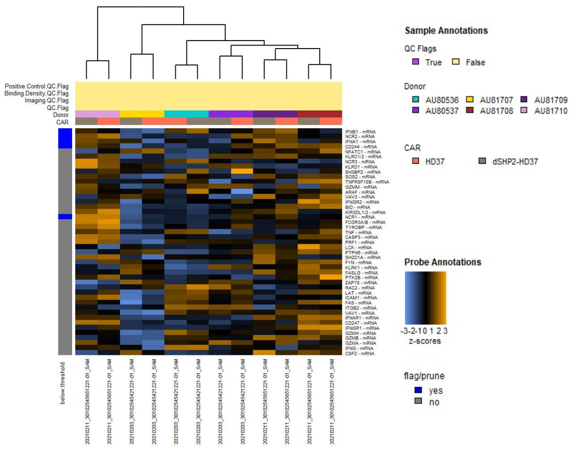


B

#### Volcano Plots

Stimulated vs unstimulated CD19 CAR

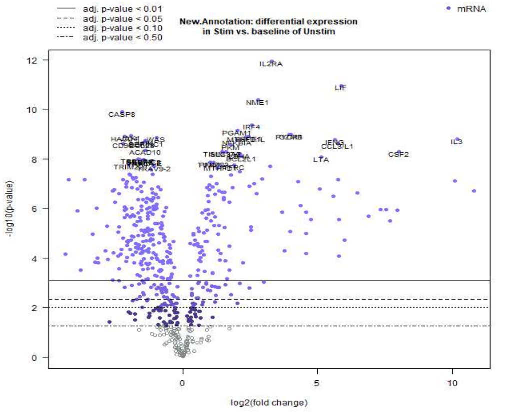


Stimulated CD19 CAR vs stimulated dSHP2 + CD19 CAR

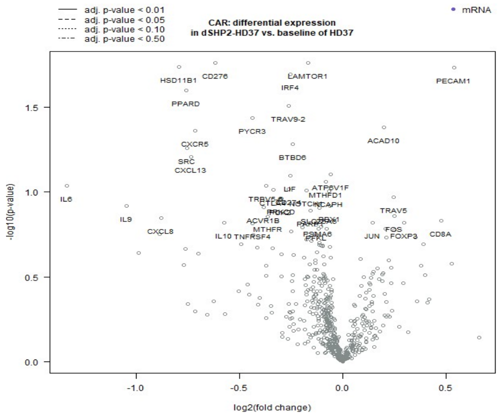


**Supplementary Figure 9: Transcriptome analysis of CAR T cells expressing dSHP.** T cells expressing the CD19 CAR in the absence or presence of dSHP2 were stimulated on plates coated with an anti-idiotype antibodies specific for the CAR binding domain. **A**) Clustering of genes associated with cytotoxicity in stimulated versus unstimulated CD19 CAR T cells (left panel) or stimulated CD19 CAR (HD37) T cells versus stimulated dSHP2 + CD19 (dSHP2-HD37) CAR T cells (right panel). **B**) Volcano plots representing differential gene expression between stimulated versus unstimulated CD19 CAR T cells (left panel) or stimulated CD19 CAR T cells versus stimulated dSHP2 + CD19 CAR T cells.

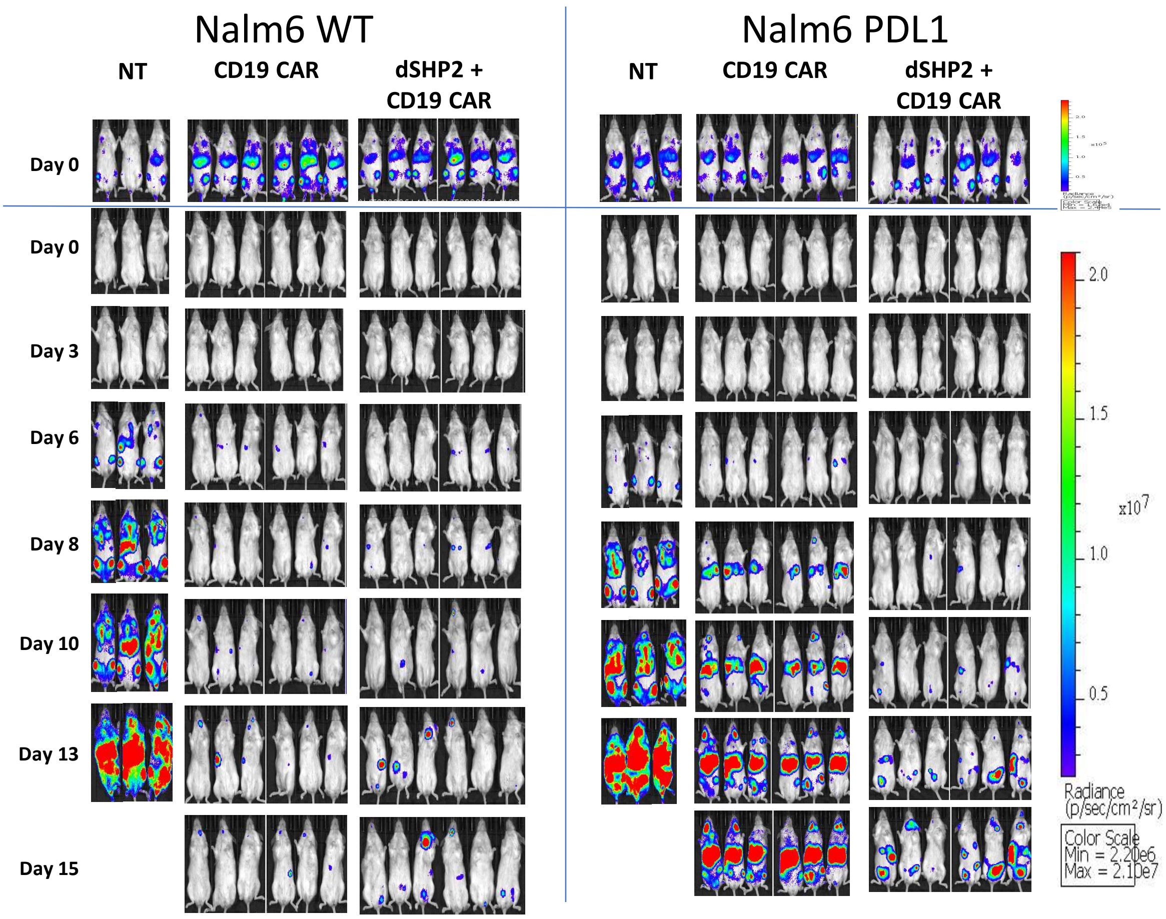


**Supplementary Figure 10: In vivo efficacy of T cells expressing CD19 CAR or dSHP2 + CD19 CAR.** Images depict data presented in Figure 4A. Images taken at Day 0 are presented with enhanced scaling (top panels) to demonstrate engraftment.

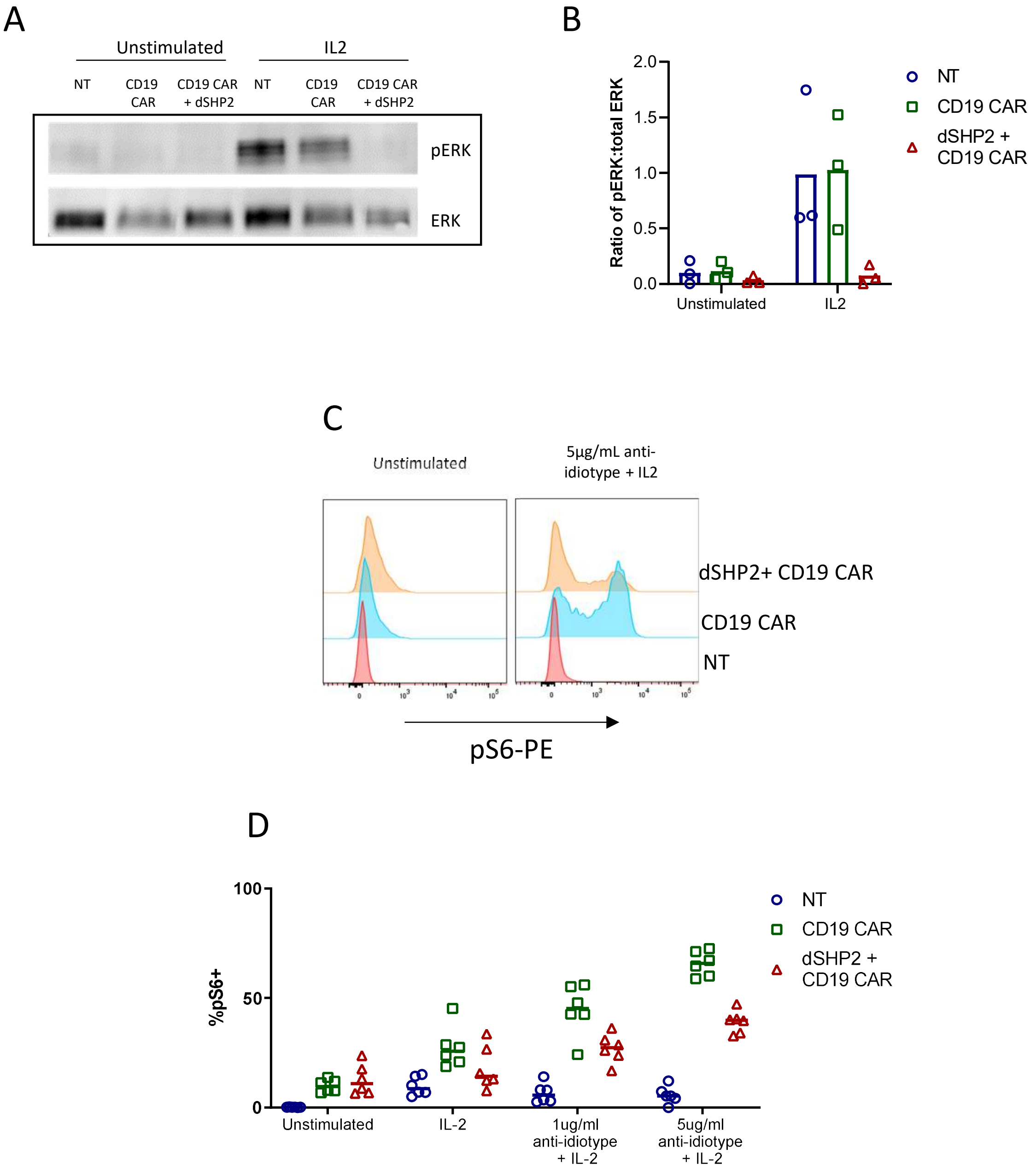


**Supplementary Figure 11: Expression of dSHP2 alters phosphorylation of Erk and ribosomal protein S6.** A) Expression of dSHP2 reduces IL2-mediated phosphorylation of ERK. T cells expressing the CD19 CAR alone or in combination with dSHP or non-transduced were stimulated with IL2 for 24 hours and phosphorylation of Erk assessed by Western blot. A representative donor is shown. B) The ratio of phosphorylated Erk: total ERK was assessed for three independent donors. C) Intracellular staining of phospho-ribosomal protein S6. T cells expressing the CAR alone or in combination with dSHP or non-transduced were left unstimulated (left panel) stimulated with CAR anti-idiotype antibody and IL2 (right panel). A representative donor is shown. D) Percentage of T cells staining for phosphorylated ribosomal protein S6 upon stimulation, as indicated, was assess for 6 independent donors.
