## Supplemental Table 1 for "Exploration of T-cell immune responses by expression of a dominant-negative SHP1 and SHP2"

| Protein name | Peptide sequence | Peptide name |
|---|---|---|
| **PD1** | KEDPSAVPVFSVDYGELDFQWRE | ITIM |
| | KEDPSAVPVFSVD**Y**GELDFQWRE | Phospho-ITIM |
| | KTPEPPVPSVPEQTEYATIVFPSGMGTSS | ITSM |
| | KTPEPPVPSVPEQTE**Y**ATIVFPSGMGTSS | Phospho-ITSM |
| **TIGIT** | LHDYFNVLSYRSLGNCSFFTETG | 1 |
| | LHD**Y**FNVLS**Y**RSLGNCSFFTETG | p1 |
| | LHD**Y**FNVLSYRSLGNCSFFTETG | p1.1 |
| | LHDYFNVLS**Y**RSLGNCSFFTETG | p1.2 |
| **2B4** | TSPKEFLTIYEDVKDLKTRRNHE | 2 |
| | TSPKEFLTI**Y**EDVKDLKTRRNHE | p2 |
| | TFPGGGSTIYSMIQSQSSAPTSQ | 3 |
| | TFPGGGSTI**Y**SMIQSQSSAPTSQ | p3 |
| | TSQEPAYTLYSLIQPSRKSGSRK | 4 |
| | TSQEPAYTL**Y**SLIQPSRKSGSRK | p4 |
| | HSPSFNSTIYEVIGKSQPKAQNP | 5 |
| | HSPSFNSTI**Y**EVIGKSQPKAQNP | p5 |
| **BTLA** | LEENKPGIVYASLNHSVIGPNSR | 6 |
| | LEENKPGIV**Y**ASLNHSVIGPNSR | p6 |
| | PNSRLARNVKEAPTEYASICVRS | 7 |
| | PNSRLARNVKEAPTE**Y**ASICVRS | p7 |
| **PAG1** | QENMVEDCLYETVKEIKEVAAAA | 12 |
| | QENMVEDCL**Y**ETVKEIKEVAAAA | p12 |
| **CTLA4** | KRSPLTTGVYVKMPPTEPECEKQ | 18 |
| | KRSPLTTGV**Y**VKMPPTEPECEKQ | p18 |
| | YVKMPPTEPECEKQFQPYFIPIN | 19 |
| | **Y**VKMPPTEPECEKQFQP**Y**FIPIN | p19 |
| | **Y**VKMPPTEPECEKQFQPYFIPIN | p19.1 |
| | YVKMPPTEPECEKQFQP**Y**FIPIN | p19.2 |

**Supplementary Table 1:** List of peptide sequences derived from the endodomains of immune inhibitory receptors. Phosphorylated tyrosines are indicated in red.
